# Characterisation of emergent toxigenic M1_UK_ *Streptococcus pyogenes* and associated sublineages

**DOI:** 10.1101/2022.12.27.522030

**Authors:** Ho Kwong Li, Xiangyun Zhi, Ana Vieira, Harry J Whitwell, Amelia Schricker, Elita Jauneikaite, Hanqi Li, Ahmed Yosef, Ivan Andrew, Laurence Game, Claire E. Turner, Theresa Lamagni, Juliana Coelho, Shiranee Sriskandan

**Affiliations:** Department of Infectious Disease, Imperial College London, UK; MRC Centre for Molecular Bacteriology & Infection (CMBI), Imperial College London, UK; National Phenome Centre and Imperial Clinical Phenotyping Centre, Department of Metabolism, Digestion and Reproduction, Imperial College London, UK; Section of Bioanalytical Chemistry, Division of Systems Medicine, Department of Metabolism, Digestion and Reproduction, Imperial College London, UK; NIHR Health Protection Unit in Healthcare-associated Infection and Antimicrobial resistance, Imperial College London, UK; School of Public Health, Imperial College London, UK; Genomics Facility, UKRI-MRC London Institute for Medical Sciences (LMS), Imperial College London, UK; The Florey Institute, School of Biosciences, University of Sheffield, UK; Centre for Infections, UK Health Security Agency, London, UK

**Author notes:** contributed equally (Ho Kwong Li, Xiangyun Zhi).

## Abstract

*Emm*1 *Streptococcus pyogenes* is a successful, globally-distributed epidemic clone that is regarded as inherently invasive. An *emm*1 sublineage, M1_UK_, that expresses increased SpeA toxin, was associated with increased scarlet fever and invasive infections in England in 2015/2016. Defined by 27 SNPs in the core genome, M1_UK_ is now dominant in England. To more fully characterise M1_UK_, we undertook comparative transcriptomic and proteomic analyses of M1_UK_ and contemporary non-M1_UK_ *emm*1 strains (M1_global_).

Just seven genes were differentially expressed by M1_UK_ compared with contemporary M1_global_ strains. In addition to speA, five genes in the operon that includes glycerol dehydrogenase were upregulated in M1_UK_ (gldA, mipB/talC, pflD, and pts system IIC and IIB components), while aquaporin (glpF2) was downregulated. M1_UK_ strains have a stop codon in gldA. Deletion of the gldA gene in M1_global_ abrogated glycerol dehydrogenase activity, and recapitulated upregulation of gene expression within the operon that includes gldA, consistent with a feedback effect.

Phylogenetic analysis identified two intermediate *emm*1 sublineages in England comprising 13/27 (M1_13SNPs_) and 23/27 SNPs (M1_23SNPs_) respectively, that had failed to expand in the population. Proteomic analysis of these four major phylogenetic *emm*1 groups highlighted sublineage-specific changes in carbohydrate metabolism, protein synthesis and protein processing; upregulation of SpeA was not observed in chemically-defined medium. In rich broth however, transcription and secretion of SpeA was upregulated ~10-fold in both M1_23SNPs_ and M1_UK_ sublineages, compared with M1_13SNPs_ and M1_global_.

We conclude that stepwise accumulation of SNPs led to the emergence of M1_UK_. While increased expression of SpeA is a key indicator of M1_UK_ and undoubtedly important, M1_UK_ strains have outcompeted M1_23SNPs_ and other *emm* types that produce similar or more superantigen toxin. We speculate that an accumulation of adaptive SNPs has contributed to a wider fitness advantage in M1_UK_ on an inherently successful *emm*1 streptococcal background.

**Data availability:** RNAseq. All new RNAseq data are uploaded to the European Nucleotide Archive under project reference PRJEB58303

Genomic data. All genomes listed are available on the European Nucleotide Archive using accession numbers as listed in the appendix,

Proteomes. Proteomic data are available on FigShare 10.6084/m9.figshare.21777809 and will be uploaded to PRIDE

**Impact Summary:** Although the major *Streptococcus pyogenes* reservoir is in children with pharyngitis and skin infections, *S. pyogenes* can lead to rarer, invasive infections that are rapidly progressive and associated with high mortality and morbidity. *Emm*1 *S. pyogenes* strains are the single most frequent genotype to cause invasive infections in high income countries and are established worldwide as an epidemic clone. The M1_UK_ *S. pyogenes emm*1 sublineage which is defined by 27 new SNPs in the core genome, and characterised by increased scarlet fever toxin SpeA production, emerged and rose to dominance over a period of 5-6 years since initial recognition, outcompeting other *emm*1 strains in England. Increased dominance of *emm*1 among invasive infections this winter, on a background of already-increased numbers of *S. pyogenes* infections, points to a key shift in host-pathogen interaction. We hypothesize that a combination of pathogen fitness, virulence, and host susceptibility have coalesced to account for the excess of circulating *S. pyogenes* and *emm*1 invasive infections. In this paper we undertake a systems-based evaluation of M1_UK_ in comparison to older non-M1_UK_ *emm*1 strains, and identify a number of pathways that are altered in addition to the previously-reported increased SpeA expression. The emergence of a new sublineage within an already virulent clone requires ongoing surveillance, and more detailed investigation of the likely mechanisms leading to increased fitness. The capacity of *S. pyogenes* to cause outbreaks at national scale highlights a potential need to consider strain-specific public health guidance, underlining the inherent virulence of this exclusively human pathogen.

## Introduction

*Emm1 Streptococcus pyogenes* emerged in the 1980’s and spread globally to become the leading cause of invasive *S. pyogenes* infection throughout the developed world (1,2). The lineage expanded following a recombination event that conferred increased expression of the streptolysin O (slo/nga) toxin locus, and was associated with specific phage content, including the phage encoding a superantigen, SpeA (1). More recently, during a period of increased scarlet fever activity in England, a new sublineage of *emm*1 *S. pyogenes* (M1_UK_) was detected and found to have expanded (3). These strains were strongly associated with not only sore throats and scarlet fever, but also increases in invasive infection (3). The earliest M1_UK_ strain detected to date was in a collection of non-invasive isolates from London in 2010, while the first invasive strains were detected in England in 2012. By 2016, the M1_UK_ sublineage represented around 80% of all invasive *emm*1 isolates in England (3); this rose to 91% by end of 2020 (4). Despite differing from older *emm*1 strains by just 27 core genome SNPs, the new sublineage was characterised by a ~ten-fold increase in expression and production of the superantigen SpeA. Since 2019, the M1_UK_ lineage has been identified elsewhere in Europe and North America (5–7).

*Emm*1 strains are the single most dominant cause of invasive *S. pyogenes* infection. In this work, we set out to characterise the wider phenotype of the new sublineage M1_UK_, and to compare M1_UK_ strains with minor sublineages that appeared briefly as intermediates, although did not expand to the extent of M1_UK_. We also examined natural mutants of M1_UK_ and the minor sublineages that provide insight into the cost-benefit balance of the changes in this new highly successful group of *S. pyogenes* M1T1 strains.

## Methods

### Bacterial strains

*S. pyogenes* strains used are outlined in Supplementary Tables S1 and S2; strains stored in 20% glycerol were streaked onto Columbia blood agar (CBA) prior to broth culture. *S. pyogenes* were cultured in Todd Hewitt Broth (THB, Oxoid, UK) or chemically defined medium (CDM) comprising iron, phosphate, magnesium, manganese, sodium acetate, calcium, sodium bicarbonate, L-cysteine, bases, vitamins and amino acids, with or without different carbon sources (Supplementary Table S3) at 37°C in 5% CO_2_.

### RNA-sequencing

RNA was extracted from four different *S. pyogenes* strains from each lineage (Supplementary Table S1), cultured in THB for six hours corresponding to late-log growth phase using methods as previously described (3). RNA sequencing of M1_global_ and M1_UK_ RNA was undertaken by Novogene, Cambridge, UK and by the MRC London Institute of Medical Sciences (LMS). Data (deposited in project PRJEB58303) were analyzed according to published guidelines (8). Briefly, read quality was accessed using FastQC (https://www.bioinformatics.babraham.ac.uk/projects/fastqc/), filtered and trimmed using trimmomatic (9), and mapped against the MGAS5005 (CP000017) reference genome using bowtie2 (10) with the highest sensitivity options. The resulting alignments were converted to sorted BAM files using vcftools (11). Initial visualizations of the sequencing mapping were performed using the Integrative Genomics Viewer (IGV) (12) including confirmation of *gldA* disruption. The mapped RNA-seq reads were then transformed into a fragment count per gene per sample using HT-seq (13) package. Exploratory data analysis (Principal component analysis and Heatmap of sample-to-sample distances) of the RNAseq data was implemented and plotted using DESeq2 package (14). Differential expression analysis in each dataset was performed using three different R packages (DESeq2 (14), EdgeR (15) and limma (https://bioconductor.riken.jp/packages/3.0/bioc/html/limma.html)) with a log_2_fold change of 0.5 and p-adj < 0.05 for M1_global_ vs. M1_UK_, and a log_2_fold change of 1 and p-adj < 0.05 for M1_H1488ΔgldA_ vs. M1_H1488_. Only genes DE in two of the three softwares used were considered as DE genes and used in analysis. Prophage regions were predicted using phaster (16), and curated by visual assessment and blast alignment.

### Gene transcription studies

Specific transcript abundance was evaluated by quantitative RT-PCR using a plasmid standard for each gene and compared with *proS*. For the gldA operon plasmid standard, single amplicons were amplified to create a single linear insert (ProS-gldA-mipB-pflD-pts subunit IIC) that was TA-cloned into plasmid PCR2.1. For *glpF2* and *speA*, the plasmid standard comprised just *glpF2* and *proS*, or *speA* and *proS* respectively. cDNA synthesis from *S. pyogenes* RNA was undertaken as previously reported prior to RT-PCR (3); primers are listed in Supplementary Table S4. Comparisons were subject to analysis in GraphPad Prism v9. Non-parametric (Mann Whitney U) or t-tests were used; p<0.05 was considered significant.

### Genetic manipulation

The gene encoding gldA was mutated by allelic replacement using the suicide vector pUCMUT. A 541 bp fragment upstream of *gldA* gene was amplified (forward primer: *5’*-AGC*GAATTC*TCGCCCAAGATTACGAAGG-*3’*, reverse primer: *5’*-GG*GGTACC*CGTTGAACTCCTTTATCTGTGATT-*3’*) incorporating *5’* EcoRI and 3’KpnI restriction sites, and cloned into the suicide vector pUCMUT to produce vector pUCMUT_gldAUP_. A 532 bp fragment downstream of the *gldA* gene was amplified (forward primer: *5’*-AA*CTGCAG*CTATTGCAGAGCTGGTGCT-*3’*, reverse primer: *5’* - ACGC*GTCGAC*CGAGTCGATAGGCTAACC-*3’*) incorporating *5’* PstI and *3’* SalI restriction sites and cloned into PstI/SalI digested pUCMUT_gldAUP_ to create pUCMUTgldA_KO_. The construct was introduced into *S. pyogenes* M1_global_ strains H1488 (M1_H1488_), and BHS162 (M1_BHS162_) by electroporation and crossed into the chromosome by homologous recombination. Transformants were selected using kanamycin (400μg/ml). Successful disruption of the gldA gene and insertion of the kanamycin resistance cassette was confirmed by PCR, DNA sequencing and whole genome sequencing of mutated strains M1_H1488DgldA_ and M1_BHS162DgldA_ (isolate identifiers H1589 and H2151 respectively).

### GldA activity assay

Cell free extracts were prepared from bacteria cultured overnight in chemically defined medium containing either 0.5% glucose of 0.5% glycerol to A_600_ of 0.6-0.7 (or as close to this as feasible). Bacteria were washed, centrifuged and kept on ice for 1h within an anaerobic jar, then suspended in 10 mM Tris buffer, pH 9. cells were disrupted by agitation in three 60-second bursts with 0.1 mm glass beads. Beads were allowed to settle, and the supernatant fluid centrifuged in an Eppendorf microcentrifuge for 30 seconds at 14,000 x g. GldA results in conversion of glycerol + NAD to dihydroxyacetone + NADH +H^+^. GldA activity was derived from the increase in absorbance at 340 nm resulting from the reduction of NAD; one unit reduces one micromole of NAD per minute at 25°C and pH 10.0 under the conditions specific (17).

### Phylogenetic analysis

*Emm1* genomes used in phylogenetic analysis were from the UK and are listed in the Supplementary Information. These comprise sequenced non-invasive *emm*1 isolates (n=139) (3); sequenced invasive *emm*1 isolates (n=40) from two studies (3,18); 64 invasive *emm*1 isolates from the British Society for Antimicrobial Chemotherapy (BSAC) collection (19); and 23 *emm*1 isolates from a hospital outbreak study (20). Two new *emm*1 genomes were sequenced from an additional outbreak and are available from the European nucleotide archive (Project PRJEB36425: ERS4267588 and ERS4267589). Raw reads were trimmed using trimmomatic version 0.36 (9) with the default parameters. The SNP calling was performed by mapping trimmed reads to the complete *emm*1.0 MGAS5005 (CP000017) reference genome using Snippy v4.6.0 (https://github.com/tseemann/snippy), with a minimum coverage of 10, minimum fraction of 0.9, and minimum vcf variant call quality of 100. Gubbins version 2.4.1 (21) was used to identify and remove recombinant regions from the resulting full genome alignment file. A maximum likelihood phylogeny was created from core SNPs using the general time-reversible (GTR) model of nucleotide substitution with the gamma distributed rate heterogeneity implemented in FastTree v2.1.10-4 (22) Phylogenetic trees were visualized using FigTree v1.4.2 (http://tree.bio.ed.ac.uk/software/figtree/) and Microreact (https://microreact.org/showcase) and edited using INKSCAPE (https://inkscape.org/pt/).

### SpeA expression

Semi-quantitative analysis of SpeA expression by *S. pyogenes* cultured in THB for 16 hours, was undertaken using cell-free culture supernatants concentrated 5X using Amicon filters, western blotting using a rabbit polyclonal antibody to SpeA and comparison with standard concentrations of rSpeA expressed from *Escherichia coli* as previously reported (3).

### Proteomics

In pilot studies, five strains were randomly selected from each of M1_UK_ or M1_global_, cultured in 50mL chemically defined medium (CDM) to A_600_ 1.2-1.4 (6 hours) at 37°C with 5% CO_2_, then cytosolic, cell wall, and supernatant fractions prepared for proteomic analysis. For proteomic analysis of sublineages, strains were randomly selected from five phylogenetic branches within each lineage (5 strains per sublineage, 4 sublineages in total). The supernatant fraction was removed, syringe filtered (Minisart 0.2uM filter, Sartorius, Germany) and proteins precipitated overnight at 4°C using 10% Trichloroacetic acid precipitation. Cell-wall-proteins were extracted from the bacterial pellet using 1mL of 30% raffinose, centrifugation at 10,000 RPM for 5 min, followed by resuspending the pellet in 1mL of cell wall extraction buffer (960μL 30% Raffinose, 10μL 1M Tris-HCl pH8, 10μL of 10kU/mL mutanolysin, 10μL 100mg/mL lysozyme, and 10μL protease inhibitor cocktail III (Avantor VWR, USA), followed by incubation at 37°C for 3 hours with occasional turning, and then aspiration of cell wall extract supernatant after centrifugation at 13,000 RPM for 10 minutes. The residual cytosolic fraction was further mechanically lysed via bead beating for 3 cycles for 45 seconds (Lysing Matrix B from MP Bio, USA). The samples of each cellular fraction then underwent centrifugal concentration using 3kDa filters and buffer exchanged (Amicon Ultra-15, Millipore, USA) with 50mM Tris buffer at pH8. The samples were then submitted to the Proteomics Facility of the National Phenome Centre (London, UK) for LC MSe (Data to be deposited in PRIDE; currently deposited in FigShare). Precipitated samples were dissolved in 8M urea, 100mM ammonium bicarbonate (AmBic) by sonicating for 10 minutes in a water bath. Total protein was determined in all samples by Protein Assay (Protein Assay II, BioRad) according to the manufacturer’s instructions. 20μg of protein was digested by the addition of 40mM chloracetamide, 10mM TCEP (Bondbreaker, ThermoScientific) and 0.2μg of trypsin in 100mM AmBic. Proteins in 8M urea were diluted to 1M urea prior to the addition of trypsin and all samples left overnight at 37°C. Desalting was performed by acidifying samples to 0.5% trifluroacetic acid (TFA) and adding them to a pre-equilibrated uElution HLB desalting plate (Waters), washing (3×100μl) with 0.5% TFA and eluding with 80% acetonitrile (3×50μl). All washes were drawn through the plate under vacuum. Desalted peptides were dried completely at 45°C in a vacuum-centrifuge.

For mass-spectrometry analysis, proteins were dissolved in 0.1% formic acid by sonicating in a water bath for 10 minutes. 0.5μg of peptides were analysed by LC-HDMSE (M-class UHPLC (Waters), Synaptic G2S (Waters)). Data was searched and processed using Progenesis QI for Proteomics.

Differentially expressed proteins with a fold change threshold of log_2_ 1.5 (p value threshold 0.05) were visualized on volcano plots. Enrichment analysis and protein-protein interactions were performed using STRING (https://string-db.org/), a database able to predict direct (physical) and indirect (functional) associations based on collected data across a range of experimental and *in silico* protein interactions. Proteins with a percentage identity higher than 90% and a “combined interaction score” higher than 0.7 were used to create a protein network in which the interaction between two proteins was inferred based on the information available in the STRING database and colour coded accordingly.

## Results

### Transcriptome of M1_UK_ S. pyogenes

When comparing broth-cultured M1_UK_ and M1_global_ *S. pyogenes*,significant differential expression of just seven genes was observed (Table 1). As expected, transcription of SpeA was upregulated in all M1_UK_ strains compared with other M1_global_ strains; increased speA transcription by M1_UK_ has previously been confirmed by RT-qPCR of RNA from 135 *emm*1 isolates (3). Unexpectedly, transcription of glpF2, a putative aquaporin (identified as Spy1573 in *emm*1 reference strain MGAS5005), was markedly downregulated in M1_UK_ strains. Bioinformatic analysis of the *S. pyogenes* aquaporin gene glpF2 demonstrated similarity to the glpF3 family of *Lactobacillus plantarum* reported to be associated with both glycerol and water, but also dihydroxyacetone (DHA) transport (23).

**Table 1.** Differentially expressed genes comparing three M1_UK_ and three M1_global_ strains

| Gene ID | Gene Name | Description | Average log2foldchange | Average padj | Strand |
| --- | --- | --- | --- | --- | --- |
| M5005_Spy0996 | *speA2* | enterotoxin | 2.361 | 3.731 E-09 | + |
| M5005_Spy1573 | *glpF.2* | glycerol uptake facilitator protein | -2.423 | 5.143E-09 | - |
| M5005_Spy1742 | *mipB* | transaldolase | 1.043 | 0.0002 | - |
| M5005_Spy1741 | *gldA* | glycerol dehydrogenase | 1.024 | 0.0006 |  |
| M5005_Spy1743 | *pflD* | formate acetyltransferase | 0.963 | 0.0007 |  |
| M5005_Spy1744 | NA | PTS system, cellobiose-specific IIC component | 0.653 | 0.0061 | - |
| M5005_Spy1745 | NA | PTS system, cellobiose-specific IIB component | 0.749 | 0.0204 |  |

The remaining five differentially-expressed transcripts that were upregulated in M1_UK_ represented consecutive open reading frames in an apparent operon that includes glycerol dehydrogenase (gldA), pyruvate formate lyase (pflD), and a transaldolase-like protein (talC or mipB) as well as PTS system IIC and IIB components, annotated as cellobiose-specific.

### GldA operon

A single SNP in the glycerol dehydrogenase gene *gldA* among all M1_UK_ strains is known to introduce a premature stop codon at position 175 of the 362 residue enzyme and is predicted to result in a truncated protein with abrogated enzyme activity (3). GldA is the final open reading frame in the sequence of genes that was found to be differentially expressed (Figure 1A). Differential expression of genes comprising the apparent operon was confirmed using RT-qPCR (Figure 1B-E). Transcription of the aquaporin gene was evaluated in three strains from each lineage, and although non significant, there was a 2-fold reduction in transcription in M1_UK_. (Supplementary Figure S1)

**Figure 1.**
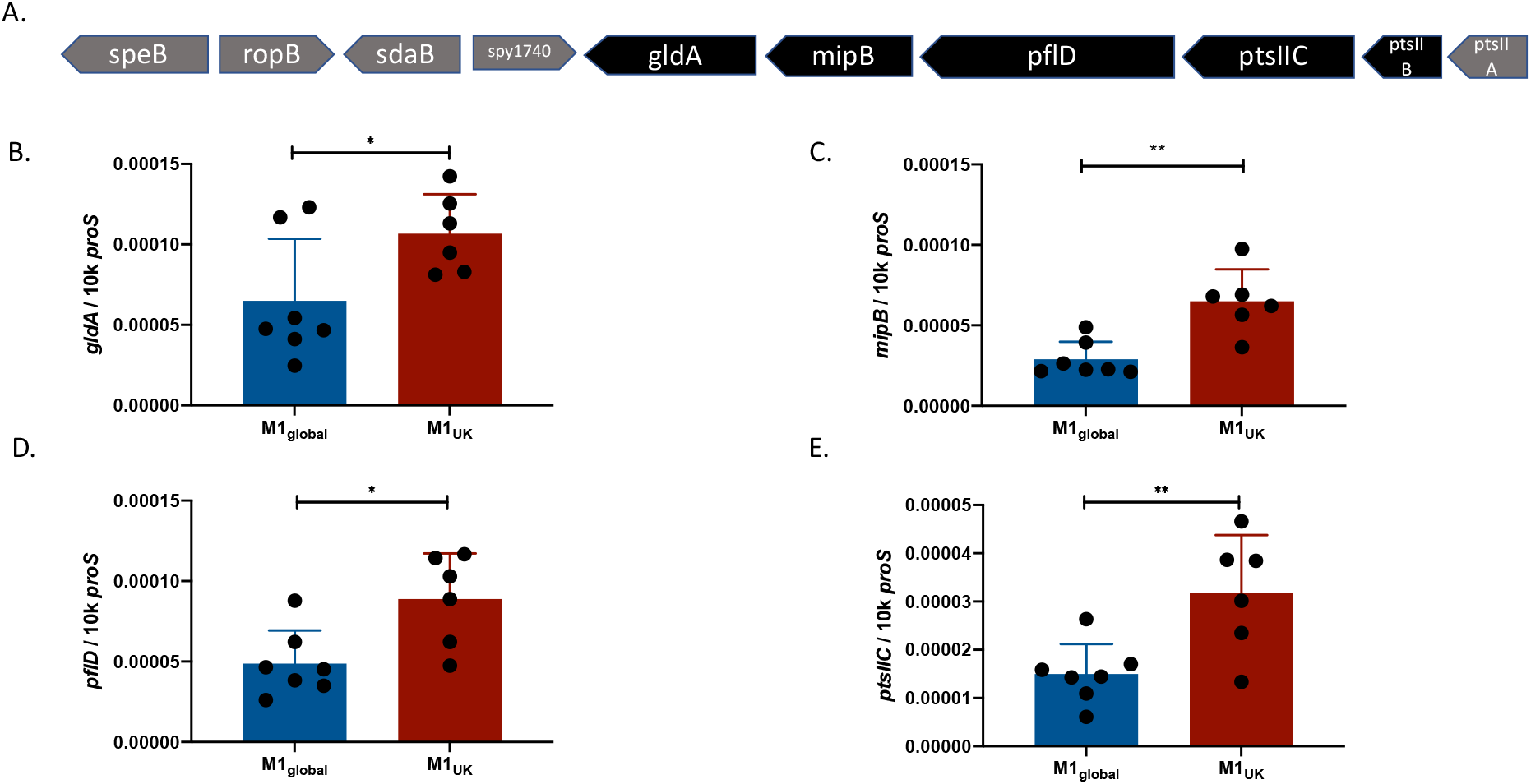
The genes within the *pflD-mipB-gldA* operon are upregulated in M1_UK_. Five adjacent genes were found to be upregulated in M1_UK_ compared to M1_global_ (A). Genes upregulated in RNAseq are shown in black and include two components of a PTS system annotated as a PTS system (cellobiose) subunits IIC and IIB. Quantitative real time PCR using RNA from M1_global_ (n=7) and M1_UK_ (n=6) strains indicating transcription of gldA (B); mipB, also known as talC (C); pflD (D); and PTS subunit IIC (E). Data points (black dots) represent individual strains tested as technical triplicates and expressed as copies per 10,000 copies proS. Error bars show SD of the mean.**p<0.01 using unpaired t-test; *p<0.05.

We hypothesised that the loss of GldA enzyme activity may in some way feedback on transcription of the adjacent PTS subunit EII genes, as well as *mipB* and *pflD*. To determine the impact of isolated loss of GldA function in *S. pyogenes, gldA* was disrupted through allelic replacement in M1_global_ strain M1_H1488_ to create M1_H1488ΔgldA_. A GldA enzyme activity assay was undertaken, in the presence of glycerol and glucose, to confirm that enzyme function was present in the parent strain, but abrogated in the mutant (Figure 2A-B); this was replicated using a second pair of isogenic M1_global_ strains (M1_BHS162_ and M1_BHS162ΔgldA_). By comparison, M1_UK_ strain BHS581 demonstrated barely detectable gldA activity, similar to the knockouts. RNA from M1_H1488_ and the isogenic M1 _H1488ΔgldA_ was subject to RNAseq to compare the wider transcriptome of *S. pyogenes* in the absence of a functional gldA gene. Surprisingly there were almost no changes in the transcriptome except in the genes of the putative ‘gldA’ operon; deletion of *gldA* abrogated transcription of *gldA* as expected, but was associated with a clear increase in transcription of *pflD, mipB*, and the adjacent PTS system cellobiose-specific IIC genes. (Table 2). Upregulation of two adjacent genes Spy0123 and Spy0124 (including sloR) was also observed.

**Figure 2.**
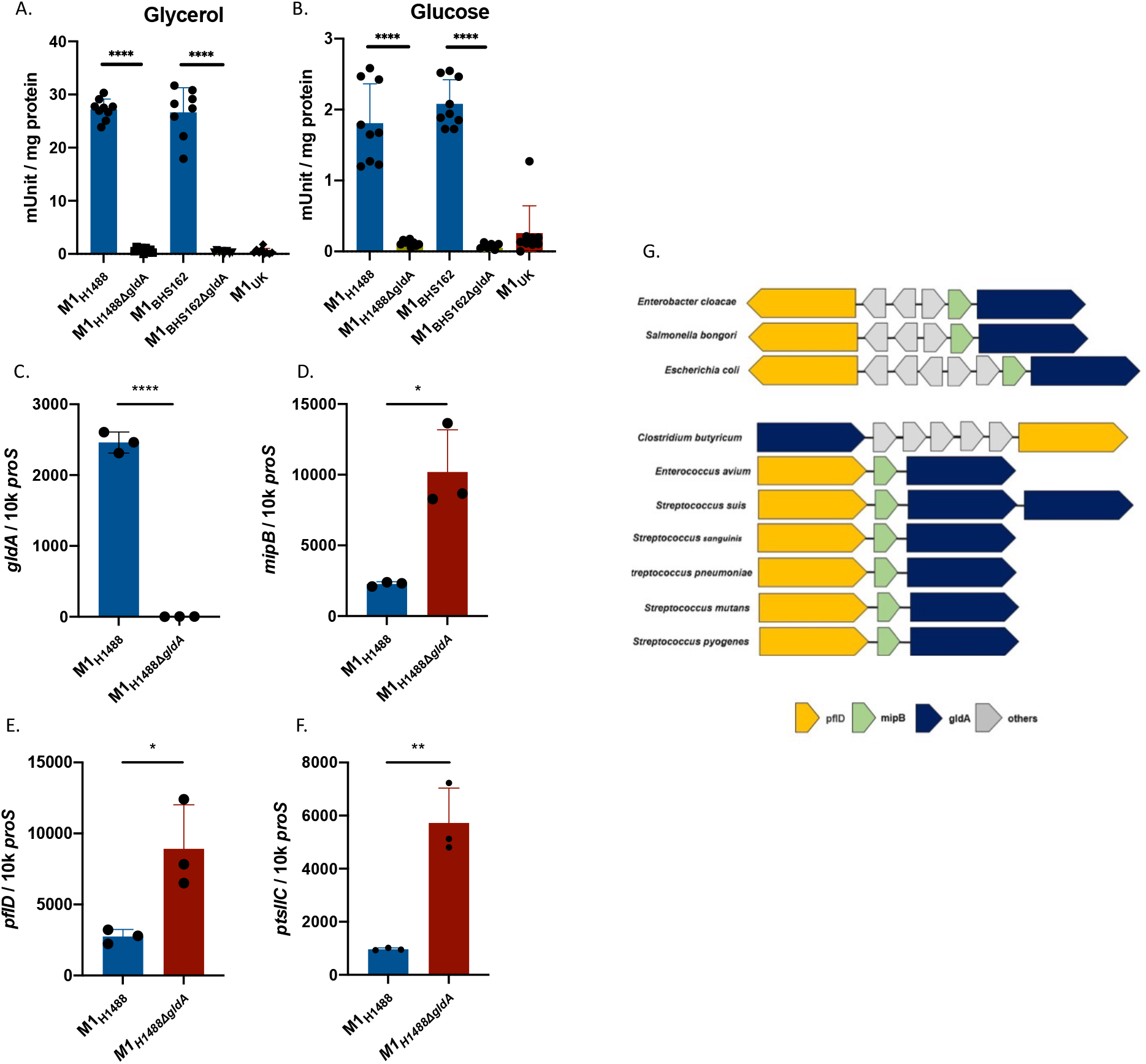
Loss of gldA function is associated with upregulated expression of adjacent genes. Glycerol dehydrogenase activity in *S. pyogenes* M1_global_ is abrogated following inactivation of the gldA gene, to the level observed in M1_UK_. Activity in the parent strain was greatest when cultured in the presence of glycerol (A) and less in glucose (B). Data show 8 or 9 individual reactions for each strain. Deletion of the gldA gene resulted in markedly reduced expression of gldA, (C), upregulated transcription of mipB (D); pflD (E); and PTS component IIC (F). Data show three biological replicates per strain. Error bars show SD of the mean *p<0.05; **p<0.01; ****p<0.0001 using Mann-Whitney U (A-B) & unpaired t-test (C-F). (G) Arrangement of pflD, mipB, gldA genes in different bacterial species.

**Table 2.**
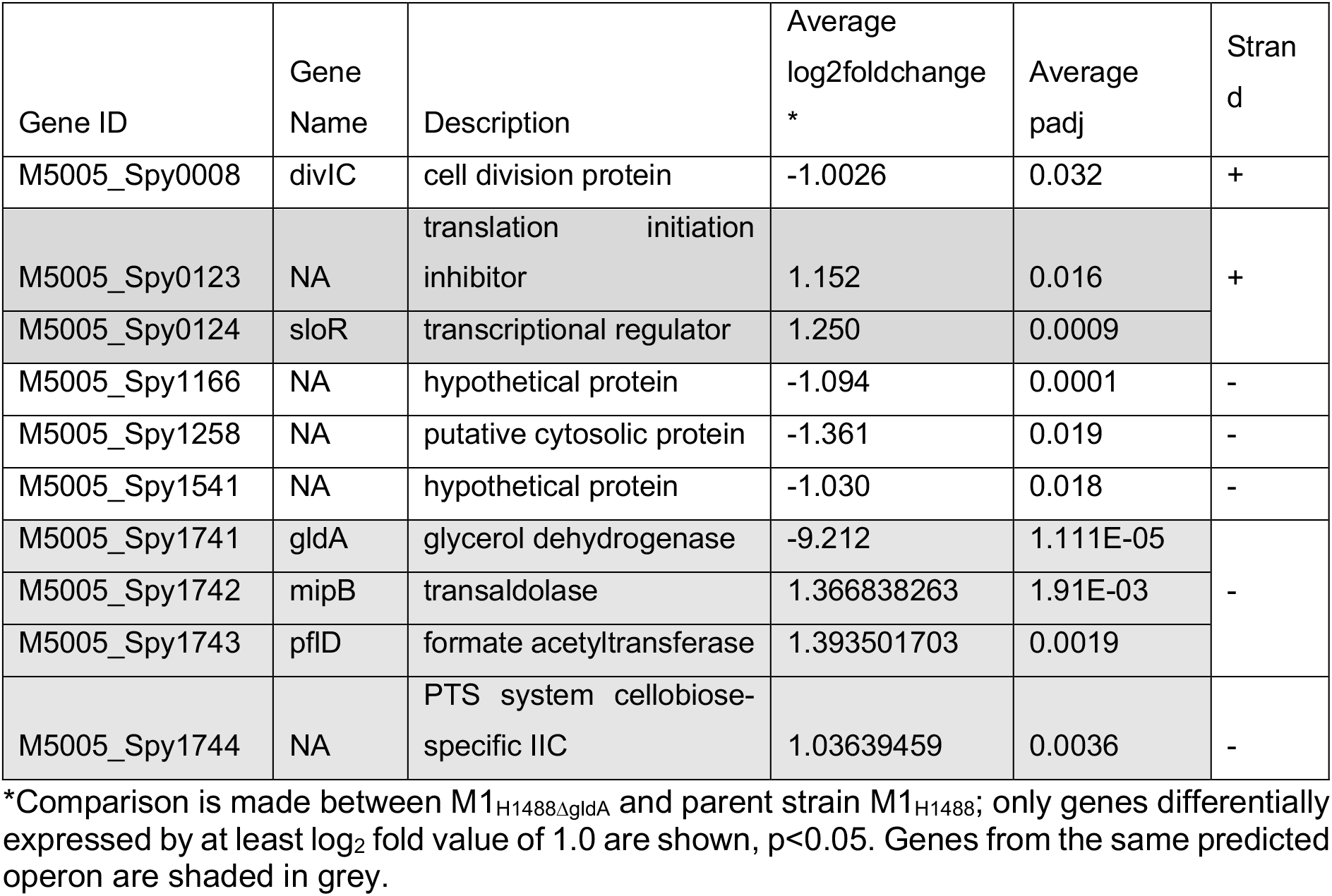
RNAseq comparison of gldA-mutant *S. pyogenes* and parent strain

Significant downregulation of gldA transcription, and upregulation of the adjacent genes was confirmed by RTqPCR (Figure 2 C-F). Taken together, the data suggested that loss of gldA activity led to upregulation of the entire operon that is concerned with metabolism of dihydroxyacetone (DHA), fructose 1, 6, phosphate, and pyruvate. *S. pyogenes* has been reported to use a number of carbon sources, however, under conditions where *emm1 S. pyogenes* grew well in CDM supplemented with glucose, we were unable to demonstrate any growth in CDM supplemented with glycerol alone, consistent with other reports (24) (not shown). Informatic analysis of publicly available genomes from a range of bacterial species demonstrated remarkable conservation of the genes and organisation of this region in all members of the streptococcaciae. Whereas other bacterial species possessed the three genes mipB (also annotated as talC, for transaldolase), pflD and gldA, the organisation of genes differed widely (Figure 2G).

### Intermediate sublineages of emm1 S. pyogenes

M1_UK_ strains are distinguished from older *emm*1 strains by the presence of 27SNPs (3) (Table 3). Although a number of additional indels are common in M1_UK_, only the 27SNPs define the new lineage. When analysing genomes from *S. pyogenes* strains isolated in the United Kingdom, we identified small numbers of strains with either 13 of the 27SNPs, or 23 of the 27SNPs (3). All *emm*1 sublineages bar M1_global_ possessed three SNPs in the transcriptional regulator RofA, however the gldA stop codon is present only in strains with 23SNPs or 27SNPs. (Table 3). We analysed our original non-invasive *S. pyogenes* WGS alongside other sequenced UK *emm*1 strains (Supplementary Table 2) and enriched for sublineages by including 10 invasive isolates from each of the following groups; M1_global_; M1_13SNPs_; M1_23SNPs_; M1_UK_. (Figure 3). As reported before, the earliest M1_UK_ strain identified was 2010, however the earliest M1_13SNPs_ strain was 2005, from the BSAC collection.

**Figure 3.**
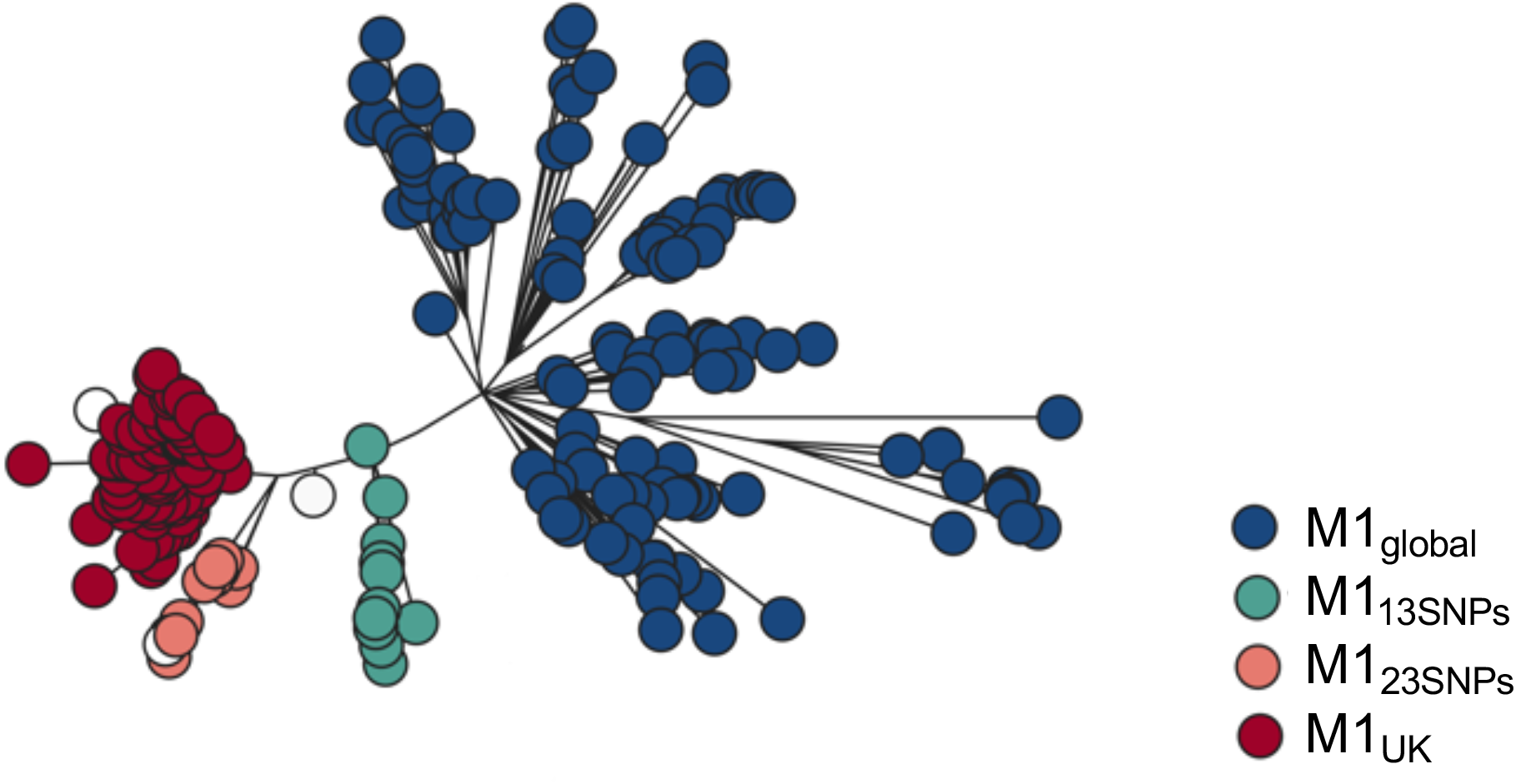
M1_UK_, M1_global_ and two intermediate sublineages Maximum likelihood phylogenetic tree constructed from core single-nucleotide polymorphisms (without recombination regions) of 269 invasive and non-invasive *emm*1 *S. pyogenes* strains representative of four main groups (M1_global_, M1_13SNPs_, M1_23SNPs_, M1_UK_). The phylogenetic tree is coloured as described in the legend. White bubbles represent isogenic strains from two distinct outbreaks with 26 and 22 SNPs, respectively and one invasive strain with 19 SNPs. Strains used in the phylogenetic tree are listed in supplementary table S2.

**Table 3.**
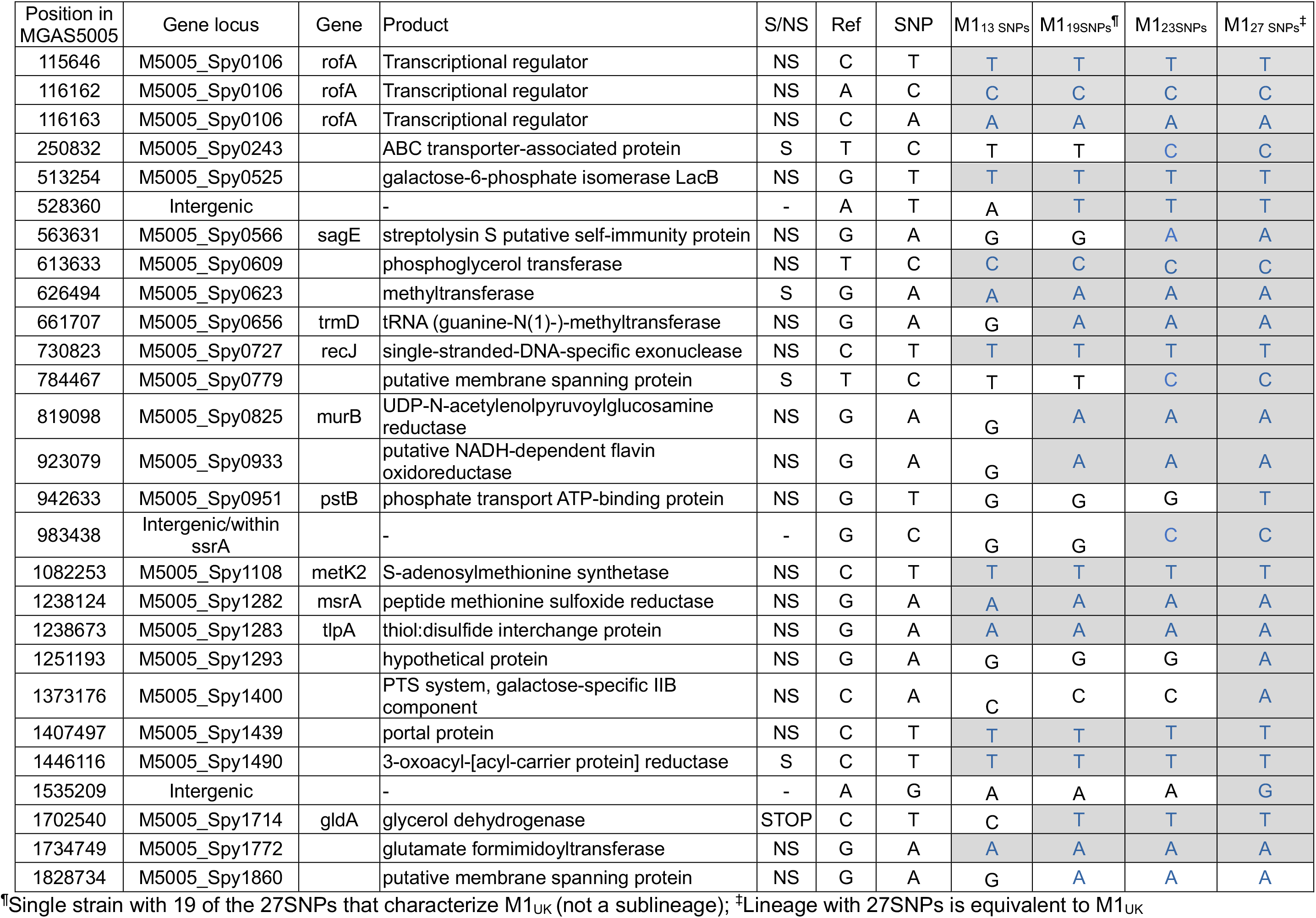
SNPs in sublineages.

| Position in MGAS5005 | Gene locus | Gene | Product | S/NS | Ref | SNP | M1<sub>13</sub> SNPs | M1<sub>19</sub>SNPs<sup>¶</sup> | M1<sub>23</sub>SNPs | M1<sub>27</sub> SNPs<sup>‡</sup> |
| --- | --- | --- | --- | --- | --- | --- | --- | --- | --- | --- |
| 115646 | M5005_Spy0106 | rofA | Transcriptional regulator | NS | C | T | T | T | T | T |
| 116162 | M5005_Spy0106 | rofA | Transcriptional regulator | NS | A | C | C | C | C | C |
| 116163 | M5005_Spy0106 | rofA | Transcriptional regulator | NS | C | A | A | A | A | A |
| 250832 | M5005_Spy0243 |  | ABC transporter-associated protein | S | T | C | T | T | C | C |
| 513254 | M5005_Spy0525 |  | galactose-6-phosphate isomerase LacB | NS | G | T | T | T | T | T |
| 528360 | Intergenic |  | - | - | A | T | A | T | T | T |
| 563631 | M5005_Spy0566 | sagE | streptolysin S putative self-immunity protein | NS | G | A | G | G | A | A |
| 613633 | M5005_Spy0609 |  | phosphoglycerol transferase | NS | T | C | C | C | C | C |
| 626494 | M5005_Spy0623 |  | methyltransferase | S | G | A | A | A | A | A |
| 661707 | M5005_Spy0656 | trmD | tRNA (guanine-N(1)-)-methyltransferase | NS | G | A | G | A | A | A |
| 730823 | M5005_Spy0727 | recJ | single-stranded-DNA-specific exonuclease | NS | C | T | T | T | T | T |
| 784467 | M5005_Spy0779 |  | putative membrane spanning protein | S | T | C | T | T | C | C |
| 819098 | M5005_Spy0825 | murB | UDP-N-acetylenolpyruvoylglucosamine reductase | NS | G | A | G | A | A | A |
| 923079 | M5005_Spy0933 |  | putative NADH-dependent flavin oxidoreductase | NS | G | A | G | A | A | A |
| 942633 | M5005_Spy0951 | pstB | phosphate transport ATP-binding protein | NS | G | T | G | G | G | T |
| 983438 | Intergenic/within ssrA |  | - | - | G | C | G | G | C | C |
| 1082253 | M5005_Spy1108 | metK2 | S-adenosylmethionine synthetase | NS | C | T | T | T | T | T |
| 1238124 | M5005_Spy1282 | msrA | peptide methionine sulfoxide reductase | NS | G | A | A | A | A | A |
| 1238673 | M5005_Spy1283 | tlpA | thiol:disulfide interchange protein | NS | G | A | A | A | A | A |
| 1251193 | M5005_Spy1293 |  | hypothetical protein | NS | G | A | G | G | G | A |
| 1373176 | M5005_Spy1400 |  | PTS system, galactose-specific IIB component | NS | C | A | C | C | C | A |
| 1407497 | M5005_Spy1439 |  | portal protein | NS | C | T | T | T | T | T |
| 1446116 | M5005_Spy1490 |  | 3-oxoacyl-[acyl-carrier protein] reductase | S | C | T | T | T | T | T |
| 1535209 | Intergenic |  | - | - | A | G | A | A | A | G |
| 1702540 | M5005_Spy1714 | gldA | glycerol dehydrogenase | STOP | C | T | C | T | T | T |
| 1734749 | M5005_Spy1772 |  | glutamate formimidoyltransferase | NS | G | A | A | A | A | A |
| 1828734 | M5005_Spy1860 |  | putative membrane spanning protein | NS | G | A | G | A | A | A |

<sup>¶</sup>Single strain with 19 of the 27SNPs that characterize M1<sub>UK</sub> (not a sublineage); <sup>‡</sup>Lineage with 27SNPs is equivalent to M1<sub>UK</sub>

### SpeA expression by sublineages

Previous comparison had demonstrated ~10-fold greater speA gene transcription by non-invasive M1_UK_ isolates compared to non-invasive M1_global_ strains (3); we first established that SpeA protein expression was similarly elevated in the same large panel of non-invasive isolates (Supplementary Figure S2). There was an indication that SpeA expression was not increased in a small number of strains from intermediate lineages. To better understand the impact of the step-wise changes in SNP content, we examined SpeA gene transcription and protein expression in a new set of strains. To include sufficient numbers of intermediate sublineage isolates, we used 40 strains from a larger national collection of invasive *emm*1 *S. pyogenes* that had been submitted to the reference laboratory and were previously sequenced (3, 18).

*SpeA* transcription was low in all M1_global_ and M1_13SNPs_ strains, except for the occasional strain with a mutation in covRS, a two component system regulator known to suppress virulence factors, but which can undergo mutation to confer a more invasive phenotype in *emm*1 and other *S. pyogenes* strains. In contrast, transcription of SpeA was high in all invasive strains with 23 or 27SNPs (Figure 4A). Likewise, SpeA protein production differed markedly between the sublineages; again SpeA production was greatest in all invasive strains with 23 or 27SNPs and was hard to detect in all M1_global_ and M1_13SNPs_ (Figure 4B). Indeed, the amount of SpeA produced routinely by M1_UK_ strains was similar to that produced by M1_global_ strains with mutations in CovRS, that is known to repress SpeA in *emm1* (25). We did not detect a difference in expression of other virulence factors such as SpyCEP, SPEB, or M protein, in broth culture (not shown). We concluded that the genetic changes required for basal increased SpeA expression in M1_UK_ resided in M1_23SNPs_ but not M1_13SNPs_.

**Figure 4.**
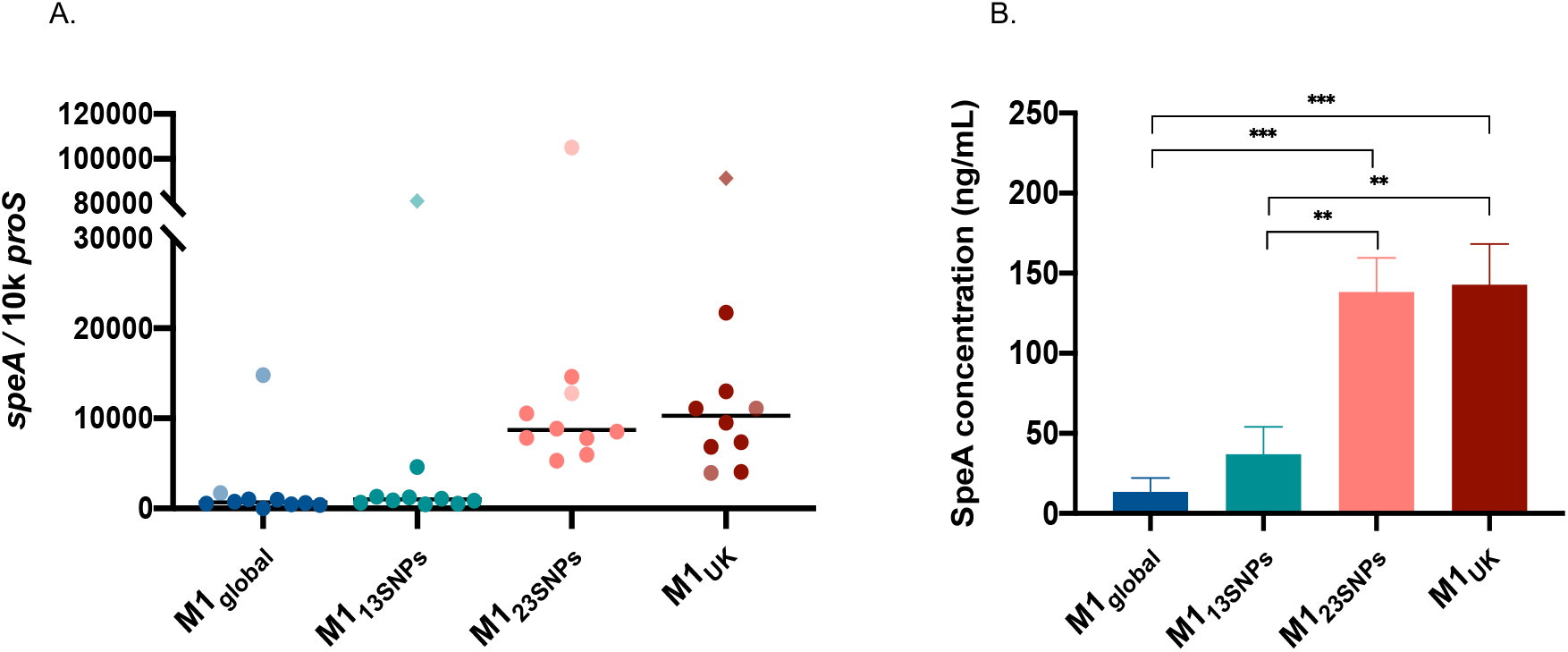
SpeA expression is increased in M1_23SNPs_ and M1_UK_ sublineages. SpeA transcription (A) using 10 strains from each sublineage is shown (total n=40). Each dot represents a single strain with lighter shading indicating the presence of *covRS* mutation (all outliers). Two isolates possess both *covRS* and *rgg4* mutations (diamond shape). Solid line represents the median. There was no statistically significant difference between the sublineages in speA transcription, largely related to the outlying covRS mutants in each sublineage. Excluding isolates with *covRS* mutation a difference was observed between M1_global_ and M1_23SNPs_ or M1_UK_ (p<0.0001), and a difference between between M1_13SNPs_ and M1_23SNPs_ or M1_UK_. (p=0.0002). SpeA protein expression (B) comprising 40 isolates (10 in each sublineage) inclusive of *covRS* mutations. Bar chart shows mean, and SEM. Multiple comparisons test made using one-way ANOVA (Tukey’s).

Isogenic isolates that differed by just single SNPs were available from two outbreak settings. Interestingly, in both settings, a single isolate was identified wherein a single SNP from the 27 SNPs that define M1_UK_ reverted to wild type. In one daycare outbreak, a non invasive isolate exhibited only 26 of the 27SNPs but was otherwise identical to an invasive isolate from the same cluster; in this case, the SNP in trmD, a tRNA (guanine-N(1)-)-methyltransferase, had reverted to wildtype. This isolate made as much SpeA as the isolate with 27SNPs.

In a separate hospital outbreak associated with a fatal case of invasive infection caused by the M1_23SNPs_ sublineage, (20), one isolate from a healthcare worker was identical to 5 other isolates in the cluster, bar one single SNP. This single SNP represented one of the 23SNPs but is present in both M1_13SNPs_ and M1_23SNPs_, a phage portal protein (Spy1439). This isolate also produced the same amount of SpeA as the parent M1_23SNPs_ strain, demonstrating the SNPs that were dispensible for increased SpeA expression.

Review of published UK *emm*1 genome sequences (19) identified a single strain with 19 of the 27SNPs among *emm*1 bloodstream isolates. Unlike the sublineage that possessed 23SNPs, this M1_19SNPs_ strain did not produce detectable quantities of SpeA, pointing to an influential role for the four SNPs that differentiate M1_19snp_ and the M1_23SNP_ sublineage in SpeA expression. Of these four SNPs, two were synonymous SNPs and felt to be unlikely to affect phenotype; one was a non-synonymous SNP in sagE; while the final change was a SNP that appeared to be intergenic in annotated *emm*1 *S. pyogenes* genomes, but lies within the start of the tmRNA ssrA (26) upstream of the phage insertion and start site of SpeA (Spy0996 in MGAS5005). RNAseq read abundance in this region did not show a difference between M1_global_ and M1_UK_ strains, with the exception of the gene encoding SpeA. Abundance of reads in the ‘paratox’ (Spy0995) gene, which is transcribed on the opposite strand to SpeA, was increased in two of four M1_UK_ strains, but this finding was not consistent.

### Proteomic analysis of S. pyogenes emm1 sublineages

To screen for lineage-specific difference in proteomes, cell wall, cytosolic, and supernatant fractions of five randomly selected M1_UK_ strains were compared with five M1_global_ following culture in CDM. Though SpeA was detected, a significant difference between M1_UK_ and M1_global_ supernatants was not observed when strains were cultured in CDM, in contrast to results (reported above) in Todd Hewitt broth, pointing to a major role for specific culture conditions in induction of SpeA. CDM supernatant from M1_UK_ strains demonstrated increased phage-encoded DNase (spd3), acid phosphatase (lppC), and a DNA binding protein. CDM supernatant from M1_global_ however demonstrated increased phosphoglycerate mutase and phosphofructokinase, both of which are linked to carbohydrate utilisation pathways in *S. pyogenes* (Figure 5A & Supplementary Figure S3A) (27). Cell wall fractions demonstrated a small number of proteins that were differentially expressed in M1_UK_ strains. These included a more than 3-fold increase in PrsA2 (Spy1732, AAZ52350.1), which controls protein folding and may operate at the ExPortal (28), and almost 2-fold increases in GAPDH and the 10kDa chaperonin groS. (Figure 5B and Supplementary Figure S3B).

**Figure 5:**
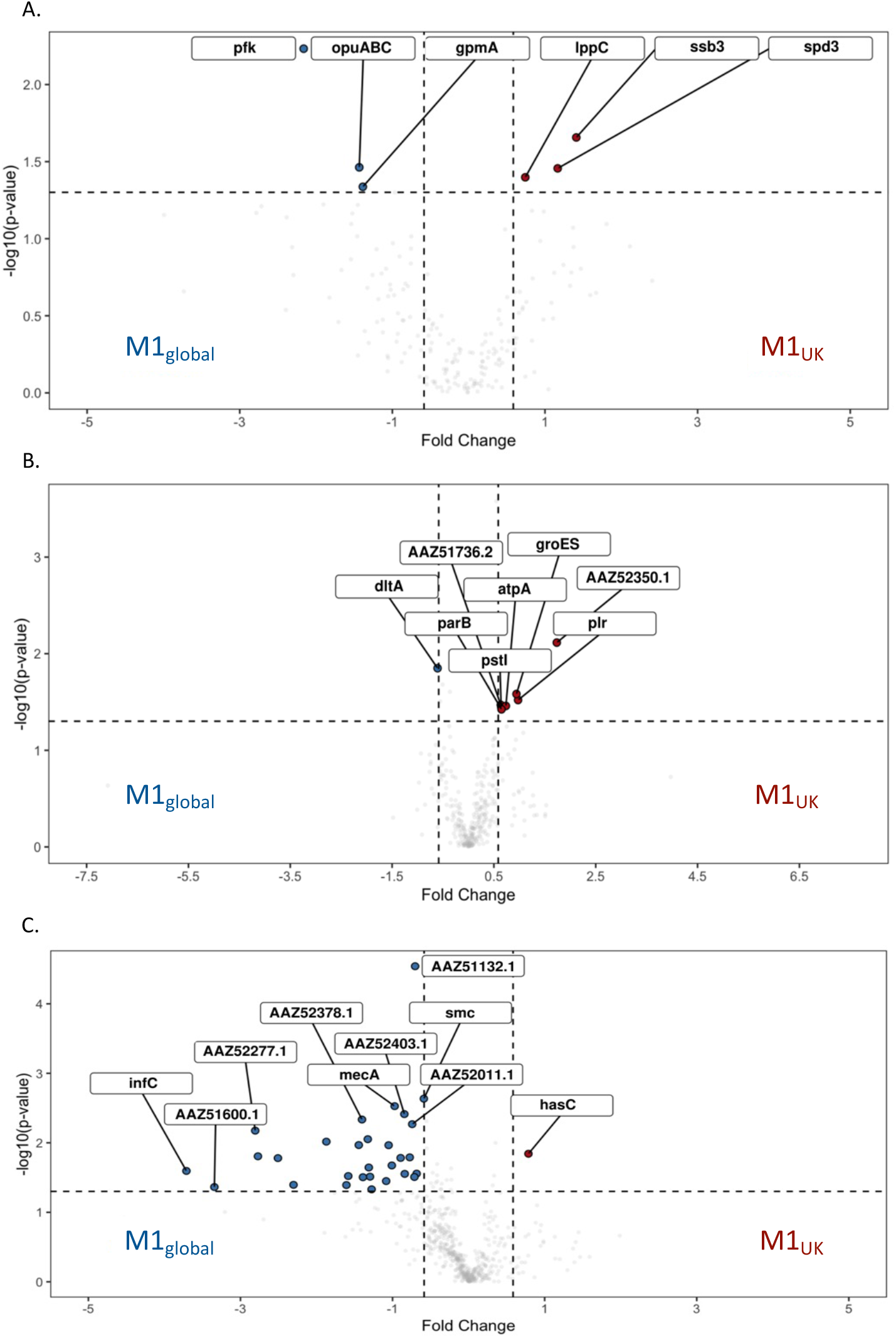
Volcano plots comparing proteins differentially expressed by M1_UK_ vs. M1_global_ cultured in CDM. Specific fractions examined were Supernatant (A); Cell wall (B); and Cytosol (C). Proteins upregulated in M1_UK_ are shown on the right in red. Those upregulated in M1_global_ are shown in the left in blue.

In M1_global_ strains, a number of cytosolic proteins were increased compared to M1_UK_, including adjacent genes Spy0438 (rnc, Ribonuclease III) and Spy0439 (smc) as well as mecA, an adapter protein and negative regulator of competence; the greatest fold changes were however seen in infC, encoding Initiation Factor 3, satD, and a number of proteins linked to protein secretion (secA), maintenance of ribosomal function and RNA. String analysis highlighted a number of links between phosphoentomutase (DeoB), protein synthesis pathways (gidA), and acid tolerance (satD). (Figure 5C and Supplementary Figure S3C).

To screen for differences between all four sublineages (two intermediate and two major sublineages), five strains from each intermediate sublineage (M1_13SNPs_ and M1_23SNPs_) were randomly selected from appropriate phylogenetic branches as well as 5 new strains from each of M1_UK_ and M1_global_. Fresh cytosolic fractions of the four phylogenetic groups (20 strains) were prepared and subject to new proteomic analysis. The data were then analysed by comparing groups in different combinations. When cytosolic preparations from all four sublineages were compared with one another, fruR expression by M1_23SNPs_ was increased in comparison to other lineages, and lowest in M1_global_, while a network of ribosomal proteins was increased in M1_13snps_ (Supplementary Figure S4A). Comparison of cytosolic preparations from new M1_UK_ and M1_global_ strains did not identify the same DE features seen previously; however a negative regulator of competence, mecA, again was increased in M1_global_ strains although only by 1.3-fold (Supplementary Figure 4B). The biggest fold change was a 3.6-fold upregulation of fruR and 5.87-fold upregulation of mur1.2 a potential autolysin (adjacent to a PTS fructose-specific IIABC system and fruR) in M1_UK_(Figure 6A). As M1_UK_ and M1_23SNPs_ strains had demonstrated comparable SpeA production, we proceeded to determine if there was commonality between these two sublineages by comparing cytosolic proteomes of [M1_UK_ and M1_23SNPs_] with [M1_global_ and M1_13SNPs_]. NtpA and B, a V type ATPase, was increased in [M1_global_ and M1_13SNPs_]; genes linked to ligase activity were found to be enriched in string analysis and highest in [M1_global_ and M1_13SNPs_](Figure 6B and Supplementary Figure S4C). When considering M1_global_ compared with all other 3 ‘new’ lineages, carbohydrate metabolism genes were further highlighted, specifically Phosphotransferase system (PTS) and disaccharide metabolic processes (Supplementary figure S4D). FruR was four-fold increased in non-M1_global_ strains, with increased FruA in M1_global_ strains; a similar pattern was seen for LacR and lacA1/lacA2 (Figure 6C, and Supplementary Figure S4D). A glutamate formiminotransferase (MGAS5005_Spy1772) was also increased in M1_global_ strains compared with non-M1_global_. Finally, comparing cytosolic proteins in M1_UK_ with all other lineages, just one protein was clearly upregulated in M1_UK_, and this was Spy0848 (ppnK), an ATP-NAD kinase. (Figure 6D and Supplementary Figure S4E).

**Figure 6:**
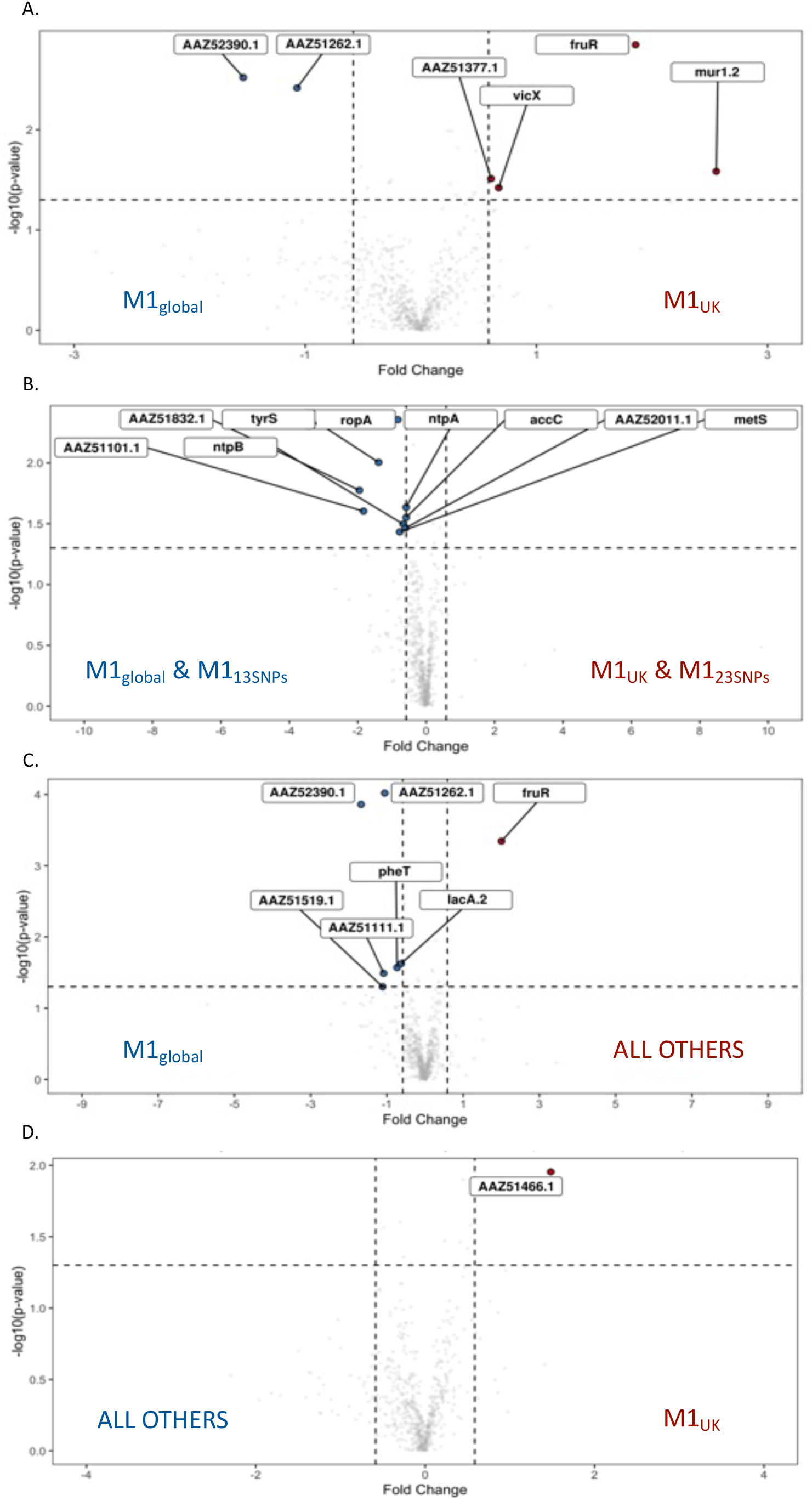
Volcano plots comparing proteins differentially expressed by different pairings of M1_global_, M1_13SNPs_, M1_23SNPs_, and M1_UK_ cultured in CDM. Cytosolic fractions only were compared as follows (A) M1_UK_ vs. M1_global_ (B) [M1_UK_ + M1_23SNPs_] vs. [M1_global_ + M1_13SNPs_] (C) All other sublineages vs. M1_global_, and (D) M1_UK_ vs. all other sublineages.

## Discussion

M1_UK_ is now the dominant *S. pyogenes emm*1 lineage in the United Kingdom, having expanded during an earlier upsurge in scarlet fever 2014-2016 (3,4). Importantly, *emm*1 strains are inherently invasive and represent the single most frequent *emm* type to cause invasive infections in the UK (29, 30). As such, any change in the *emm*1 lineage that results in increased fitness is of relevance to public health. In this first systematic study to characterize the changes in M1_UK_ and its associated lineages, we have confirmed the SpeA over-expression phenotype, and demonstrated that increased SpeA production is restricted to M1_UK_ and an increasingly rare sublineage M1_23SNPs_. The phenotype is manifest in broth culture but not CDM. M1_UK_ is defined by just 27 SNPs in the core genome including 3 SNP in rofA and a stop codon in glycerol dehydrogenase, gldA. RNA sequencing demonstrated a difference in expression of the operon that includes gldA and a phosphotransfer system (PTS) EIIC and B which represents a combined phosphate and sugar transporter, pointing to a potential shift in metabolism in the new lineage. This was accompanied by a sharp reduction in transcripts for the aquaporin gene glpF2. Preliminary proteomic analysis of strains by sublineage identified altered carbohydrate pathways related to fructose that may well be important.

Alterations in expression of gldA, mipB, pflD, and the adjacent PTS system impact on the glycolytic Embden-Meyerhof Parnas pathway (27) which, in *S. pyogenes*, relies on the phospho-enolpyruvate PTS system for acquisition of sugars other than glucose, and for transfer of phosphate ions required for carbon catabolite repression and gene regulation (31). The results indicate that both the stop codon in gldA present in M1_UK_ and allelic replacement of gldA impact glycerol dehydrogenase activity and result in upregulation of the other genes in the operon, mipB and pflD. These are involved in the glycolytic pathway required for generation and metabolism of pyruvate from glucose; the changes in carbohydrate metabolism are supported by preliminary proteomic findings that indicate alterations in fructose pathways. Interestingly, increased transcription in this operon was accompanied by increased transcription of an adjacent PTS components IIC and IIB when comparing M1_UK_ with M1_global_, and following experimental deletion of gldA. This PTS is annotated as being a cellobiose transporter; systematic experimental disruption of PTS EII systems in *S. pyogenes* has not shown a key role for these genes, however the precise sugar transported is not known (31).

The role of gldA in *S. pyogenes* has not been experimentally examined previously; gldA is reported to catalyse the conversion of glycerol to dihydroxyacetone (DHA) under microaerophilic or anaerobic conditions, however it is clear that gldA may undertake a reverse role, which is to catalyse DHA to glycerol. This may be of importance since an absence of gldA activity may lead to a build-up of DHA, which when converted to methylglyoxal can be toxic (32). The upregulation of the PTS system is of interest since these are recognised to be key players in a phosphorelay process that maintains central carbon catabolite repression of many virulence systems in *S. pyogenes* (31, 33).

The marked ~8-10 fold downregulation of aquaporin glpF2 (Spy1573) transcription, was unexpected but may represent an adaptation to the metabolic changes that have arisen in M1_UK_. There are few reports, if any, relating to glpF2 in *S. pyogenes* but there is evidence of functional links to the pflD containing operon in enterococci (34). Notably one of the intergenic SNPs that defines M1_UK_ is 39 bp from the start of the Spy1573 gene, though the significance of this is not yet known. Aquaporins are membrane proteins that function as channels for water and other uncharged solutes in all forms of life. While mostly considered as channels for water or glycerol in bacteria, potentially important to osmoregulation, aquaporins also can function as a channel for DHA. Indeed, there are similarities between glpF2 of streptococci and glpf3 of *Lactobacillus plantarum* that points to a possibility for action as a channel for DHA or similar molecules (23). Research undertaken in related *Lactococcus lactis* has also identified marked downregulation of glpF2 following osmotic stress (35). Taken together it would seem that the downregulation of glpF2 may be a necessary adaptation for M1_UK_ *S. pyogenes*, although it may also confer as-yet unknown advantage.

The upregulation of SpeA expression by M1_UK_ is clearly of importance to virulence particularly in interaction with the human host, and those who have not yet mounted an immune response to the secreted toxins of this species. There is good evidence that superantigens such as SpeA undermine development of the adaptive host immune response to *S. pyogenes* through promotion of a dysregulated T cell response associated with B cell death (36, 37). SpeA has also been shown to promote carriage of *S. pyogenes* in the nasopharynx of transgenic mice (38). To date, the expression of SpeA has only been measured in broth culture and we do not know if the upregulation in M1_UK_ might differ *in vivo*. Recent epidemiological studies found a high (44%) secondary infection rate in schoolchildren and household contacts of a case of scarlet fever caused by M1_UK_, pointing to a potential transmission advantage compared with other *S. pyogenes* lineages (39). We identified sublineage-specific altered expression of SpeA allowing us to highlight the genetic changes likely to account for this. Importantly, the three SNPs identified in the major regulator RofA do not alone account for the SpeA phenotype since these SNPs are present in M1_13SNPs_ although we cannot discount a role for these in the wider success of this lineage. While the genetic changes required for increased SpeA expression do not reside in M1_13SNPs_, they do reside in M1_23SNPs_, and strains with reversion of single SNPs pointed to a potential key role for the SNP in ssrA in SpeA upregulation. The amount of SpeA made by M1_UK_ and M1_23SNPs_ was augmented to the level of M1_global_ covRS mutants yet presumably without the fitness burden of covRS mutation that might impair pharyngeal carriage (40).

There are a number of limitations to our study. Firstly, investigation of the gldA operon is in its early stages; it is possible that the stop codon mutation in gldA confers an additional phenotype that is not recapitulated by gldA gene deletion, while the metabolic pathways that include gldA, mipB, and pflD are not fully understood. The roles of glpF2 and the PTS EII system that is upregulated are also not understood; any role in transfer of DHA for example has not been experimentally addressed. The proteomic studies are preliminary and require both validation and repetition using richer media, but have provided a rationale for further study of the role of sugar metabolism in *emm*1 *S. pyogenes*. Finally, the role of specific SNPs would necessarily require experimental proof.

Several European countries are, at the time of writing, affected by epidemic waves of invasive *S. pyogenes* disease, notably in England, where the leading cause of invasive infection is *emm*1 underlining the importance of understanding pathogenicity and transmission (30, 41). Importantly however, despite the enhanced production of SpeA by M1_23SNPs_, this intermediate sublineage did not expand in the manner seen for M1_UK_ in England, and was not detected at all in a 2020 systematic evaluation of >300 invasive *emm*1 isolates from England (4). This suggests that the fitness of M1_UK_ has required the additional acquisition of four further SNPs. These include three non-synonymous SNPs in phosphate transport ATP binding protein, pstB; a PTS galactose-specific IIB component gene; a hypothetical protein; as well as the intergenic SNP adjacent to glpF2. The amount of SpeA produced by M1_UK_ strains remains an order of magnitude lower than the amount produced by the historic *emm*1 strain NCTC8198 (42). Despite this, the new M1_UK_ lineage has outcompeted M1_23SNPs_ and has replaced older strains suggesting that the added fitness of M1_UK_ may lie beyond the ability to make SpeA.

## Supporting information

Proteomic comparisons supporting Figures 5 and 6

## Funding

This work was supported by the UK Medical Research Council (grant number MR/P022669/1); a UKRI (MRC) Research Training Fellowship (HKL); and the NIHR Imperial Biomedical Research Centre.

## Acknowledgements

SS acknowledges support from the UK National Institute for Health Research (NIHR) Health Protection Unit in Healthcare Associated Infections and Antimicrobial Resistance and the NIHR Imperial Biomedical Research Centre.

The authors acknowledge the support of colleagues in the UKRI-MRC LMS and National Phenome Centre in facilitating sequencing and proteomics.

## Disclosures

None

## Notes

### Competing Interest Statement

The authors have declared no competing interest.

