## Supplementary Figs 1-4 and Suppl. Tables 1-4 for "Characterisation of emergent toxigenic M1_UK_ *Streptococcus pyogenes* and associated sublineages"

**Supplementary file for manuscript: Characterisation of emergent toxigenic M1$_{UK}$ *Streptococcus pyogenes* and associated sublineages**

**By Ho Kwong Li*, Xiangyun Zhi*, Ana Vieira, Harry Whitwell, Amelia Schricker, Elita Jauneikaite, Hanqi Li, Ahmed Yosef, Ivan Andrew, Laurence Game, Claire E. Turner, Theresa Lamagni, Juliana Coelho, Shiranee Sriskandan**

**CONTENTS**

| | |
|---|---|
| Supplementary Figure S1 | Real-time PCR measurement of *glpF2* transcription |
| Supplementary Figure S2 | SpeA production by non-invasive *emm*1 S. pyogenes |
| Supplementary Figure S3 | String analysis of proteomic data: M1$_{global}$ vs. M1$_{UK}$ cellular fractions |
| Supplementary Figure S4 | String analysis of proteomic data: four M1 sublineages compared |
| Supplementary Table S1 | Strains used in experimental work |
| Supplementary Table S2 | Strains and WGS used for phylogenetic tree and SpeA quantification |
| Supplementary Table S3 | Chemically-defined medium composition |
| Supplementary Table S4 | Primers used for RT-qPCR |
| Excel Suppl. Files (supplied separately) | Proteomic comparisons |

**SUPPLEMENTARY FIGURES**

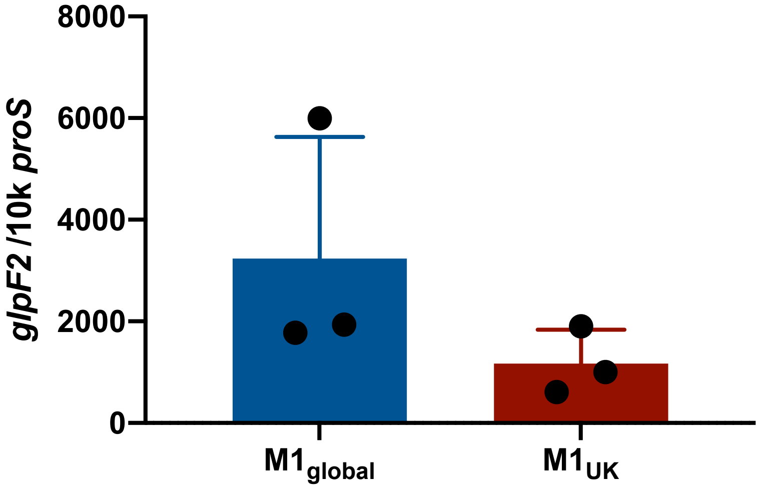

**Supplementary Figure S1**. Quantitative real-time PCR measurement of *glpF2* transcription in n=3 strains of *S. pyogenes* per lineage. Individual data points represent the average of 3 technical replicates for each strain. Strains used were: $M1_{global}$ (blue bar) (BHS0162; BHS0130; BHS0674) and $M1_{UK}$ (red bar) (BHS0258; BHS0170; BHS0128)

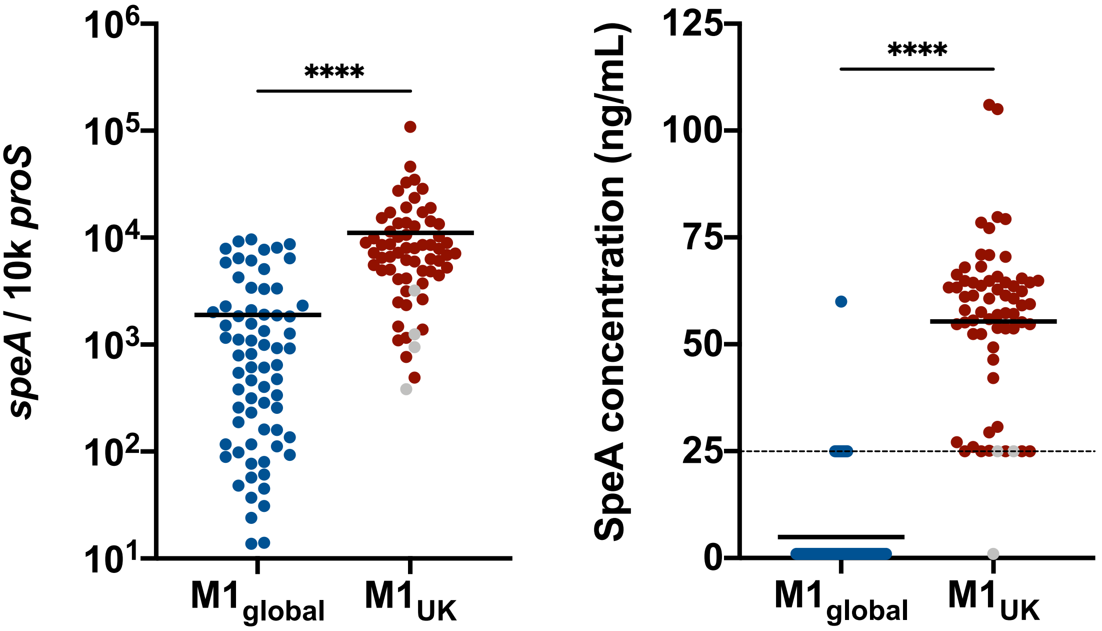

**Supplementary Figure S2**. Measurement of SpeA production by non-invasive *emm*1 *S. pyogenes* isolates, (A), Quantitative real-time PCR measurement of *speA* transcription in $M1_{global}$ and $M1_{UK}$ strains of *S. pyogenes* per lineage (previously published in Lynskey N. & Jauneikaite E. et al, Lancet Infect Dis, 2019 Nov; 19(11): 1209-1218, shown for reference only). (B) Semi quantitative measurement of SpeA expression by western blotting. N=135 strains. Solid line represents the mean, dashed line represents the limit of detection. Grey dots indicate an intermediate sublineage of M1. Unpaired T-test, **** is p<0.0001.

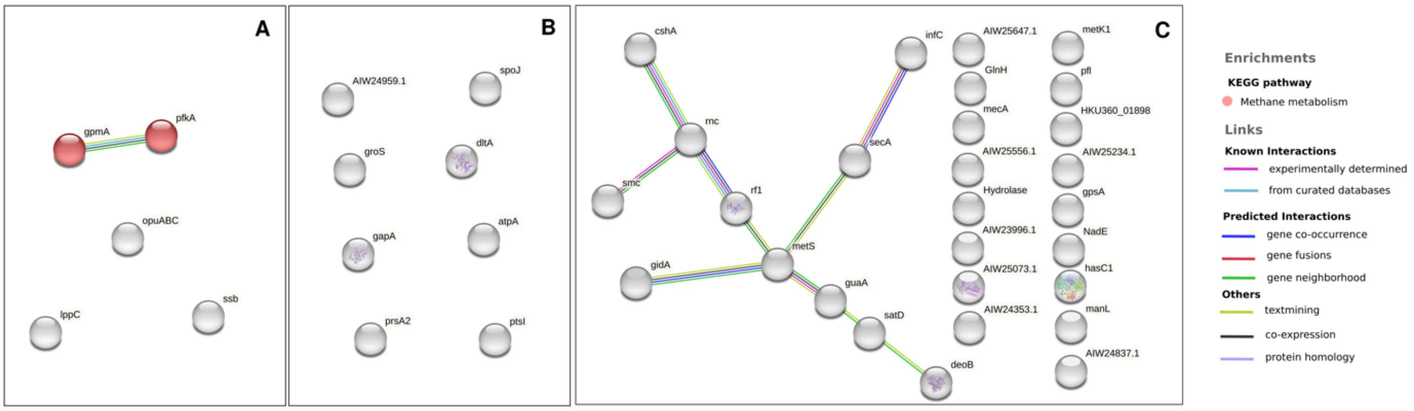

**Supplementary Figure S3. Enrichment and protein-protein interactions of differentially expressed proteins (p-value< 0.05) in different bacterial fractions using String database.** Coloured lines represent physical associations as described in the legend above. Coloured bubbles represent the type of enrichment observed. Data compare M1$_{UK}$ and M1$_{global}$ Supernatants (A); Cell wall extracts (B); Cytosol extracts (C) obtained from *S. pyogenes* strains cultured in chemically defined medium (CDM). (N=5 different strains per group).

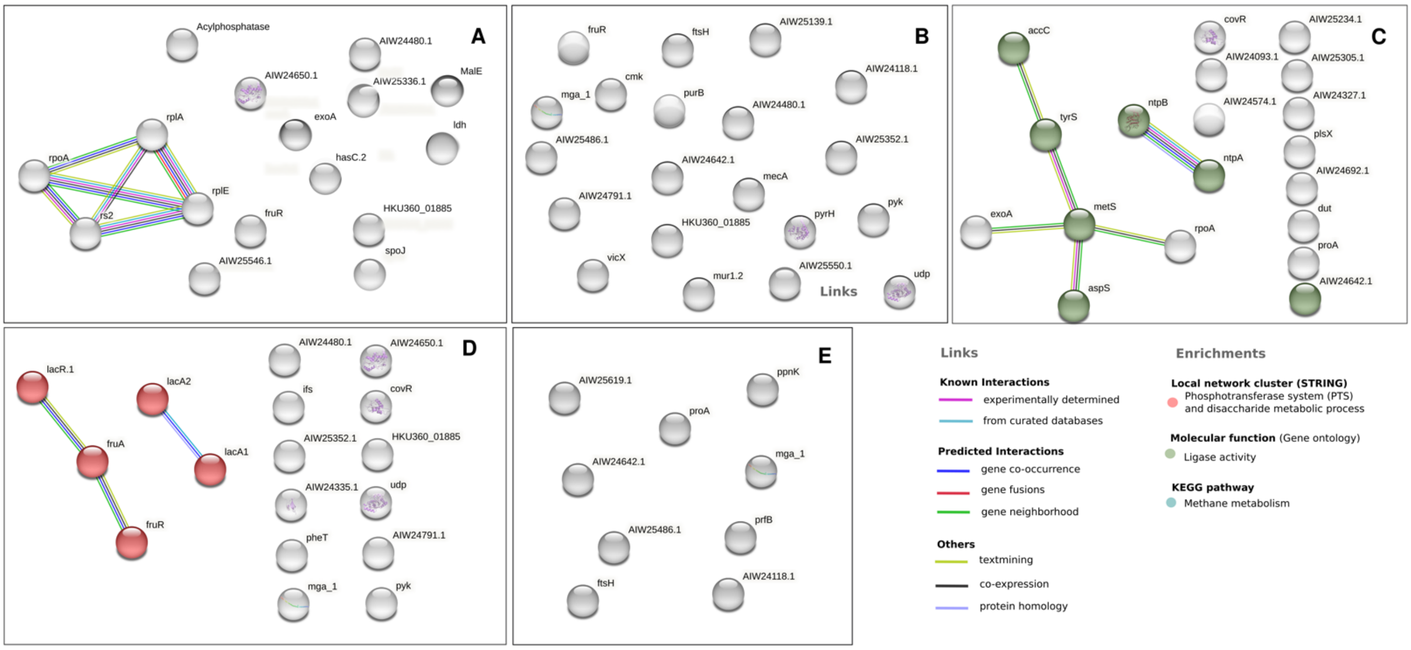

**Supplementary Figure S4. Enrichment and protein-protein interactions of differentially expressed proteins (cytosol only, p-value< 0.05) using String database comparing strains from four sublineages M1$_{UK}$; M1$_{23SNPs}$; M1$_{13SNPs}$; and M1$_{global}$ cultured in CDM** (N=5 per group). Coloured lines represent physical associations as described in the legend above. Coloured bubbles represent the type of enrichment observed. Associations shown represent: (A) All 4 groups compared, (B) M1$_{global}$ vs M1$_{UK}$, (C) [M1$_{UK}$+M1$_{23SNPs}$] vs [M1$_{13SNPs}$ + M1$_{global}$], (D) M1$_{global}$ vs all others, (E) M1$_{UK}$ vs all others.

### SUPPLEMENTARY TABLES

#### Supplementary Table S1. Bacterial strains used in RNA sequencing and RT-PCR

| | Accession¶ | Strain identifier | Lineage | Year | RNAseq | gldA operon | glpF2 |
|---|---|---|---|---|---|---|---|
| **$M1_{global}$** | ERS1020620 | BHS0674 | $M1_{global}$ | 2015 | Y | Y | Y |
| | ERS1020158 | BHS0162 | $M1_{global}$ | 2012 | Y | Y | Y |
| | ERS1020095 | BHS0130 | $M1_{global}$ | 2011 | Y | Y | Y |
| | ERS1020136 | BHS0151 or H1488 | $M1_{global}$ | 2012 | Y | | |
| | ERS1020472 | BHS0503 | $M1_{global}$ | 2010 | | Y | |
| | ERS1020045 | BHS0013 | $M1_{global}$ | 2009 | | Y | |
| | ERS1020385 | BHS0368 | $M1_{global}$ | 2009 | | Y | |
| | ERS1020523 | BHS0448 | $M1_{global}$ | 2009 | | Y | |
| **$M1_{UK}$** | ERS1020174 | BHS0170 | $M1_{UK}$ | 2012 | Y | Y | Y |
| | ERS1020090 | BHS0128 | $M1_{UK}$ | 2011 | Y | Y | Y |
| | ERS1020341 | BHS0258 | $M1_{UK}$ | 2014 | Y | Y | Y |
| | ERS1020603 | BHS0581 | $M1_{UK}$ | 2015 | Y | | |
| | ERS1020508 | BHS0521 | $M1_{UK}$ | 2015 | | Y | |
| | ERS1020714 | BHS0643 | $M1_{UK}$ | 2015 | | Y | |
| | ERS1463088 | BHS0762 | $M1_{UK}$ | 2016 | | Y | |

¶Accession numbers relate to genome sequences deposited in the European nucleotide archive

**Supplementary Table S2.** *S. pyogenes* strains and genome sequences used to create phylogenetic tree (Figure 2) and used to quantify SpeA expression by non-invasive and invasive strains.

| ENA¶† | STUDY | DISEASE | LINEAGE | LOCATION | YEAR |
|---|---|---|---|---|---|
| ERR2864947 | Sharma *et al.* | Unknown | M1global | UK | 2017 |
| ERR2864949 | Sharma *et al.* | Unknown | M1global | UK | 2017 |
| ERR2864951 | Sharma *et al.* | Unknown | M1global | UK | 2017 |
| ERR2864964 | Sharma *et al.* | Unknown | M1global | UK | 2017 |
| ERR2864950 | Sharma *et al.* | Outbreak | 23 SNPs | UK | 2017 |
| ERR2864953 | Sharma *et al.* | Outbreak | 23 SNPs | UK | 2017 |
| ERR2864957 | Sharma *et al.* | Outbreak | 22 SNPs | UK | 2017 |
| ERR2864962 | Sharma *et al.* | Outbreak | 23 SNPs | UK | 2017 |
| ERR2864966 | Sharma *et al.* | Outbreak | 23 SNPs | UK | 2017 |
| ERR2864969 | Sharma *et al.* | Outbreak | 23 SNPs | UK | 2017 |
| ERR2864948 | Sharma *et al.* | Unknown | M1uk | UK | 2017 |
| ERR2864952 | Sharma *et al.* | Unknown | M1uk | UK | 2017 |
| ERR2864954 | Sharma *et al.* | Unknown | M1uk | UK | 2017 |
| ERR2864955 | Sharma *et al.* | Unknown | M1uk | UK | 2017 |
| ERR2864956 | Sharma *et al.* | Unknown | M1uk | UK | 2017 |
| ERR2864958 | Sharma *et al.* | Unknown | M1uk | UK | 2017 |
| ERR2864959 | Sharma *et al.* | Unknown | M1uk | UK | 2017 |
| ERR2864960 | Sharma *et al.* | Unknown | M1uk | UK | 2017 |
| ERR2864961 | Sharma *et al.* | Unknown | M1uk | UK | 2017 |
| ERR2864963 | Sharma *et al.* | Unknown | M1uk | UK | 2017 |
| ERR2864965 | Sharma *et al.* | Unknown | M1uk | UK | 2017 |
| ERR2864967 | Sharma *et al.* | Unknown | M1uk | UK | 2017 |
| ERR2864968 | Sharma *et al.* | Unknown | M1uk | UK | 2017 |
| ERS4267588 | This Study | Outbreak | 26 SNPs | UK | 2018 |
| ERS4267589 | This Study | Outbreak | 27 SNPs | UK | 2018 |
| CP000017.2 | Sumby *et al.* | Reference | M1global | x | x |
| ERS1448799 | Kapatai *et al.* | Invasive | 13 SNPs | UK | 2014 |
| ERS1450651 | Kapatai *et al.* | Invasive | 13 SNPs | UK | 2015 |
| ERS1450822 | Kapatai *et al.* | Invasive | 13 SNPs | UK | 2015 |
| ERS1448193 | Kapatai *et al.* | Invasive | 23 SNPs | UK | 2014 |
| ERS1450815 | Kapatai *et al.* | Invasive | 23 SNPs | UK | 2015 |
| ERS1448173 | Kapatai *et al.* | Invasive | M1global | UK | 2014 |
| ERS1448879 | Kapatai *et al.* | Invasive | M1global | UK | 2014 |
| ERS1450607 | Kapatai *et al.* | Invasive | M1global | UK | 2015 |
| ERS1450839 | Kapatai *et al.* | Invasive | M1global | UK | 2015 |
| ERS1448481 | Kapatai *et al.* | Invasive | M1uk | UK | 2014 |
| ERS1449006 | Kapatai *et al.* | Invasive | M1uk | UK | 2014 |
| ERS1450390 | Kapatai *et al.* | Invasive | M1uk | UK | 2015 |
| ERS1450879 | Kapatai *et al.* | Invasive | M1uk | UK | 2015 |
| ERS1594714 | Lynskey & Jauneikaite *et al.* | Invasive | 13 SNPs | UK | 2013 |
| ERS1594852 | Lynskey & Jauneikaite *et al.* | Invasive | 13 SNPs | UK | 2013 |
| ERS1594863 | Lynskey & Jauneikaite *et al.* | Invasive | 13 SNPs | UK | 2013 |
| ERS1594843 | Lynskey & Jauneikaite *et al.* | Invasive | 13 SNPs | UK | 2013 |
| ERS1594990 | Lynskey & Jauneikaite *et al.* | Invasive | 13 SNPs | UK | 2016 |
| ERS1594904 | Lynskey & Jauneikaite *et al.* | Invasive | 13 SNPs | UK | 2016 |
| ERS1594914 | Lynskey & Jauneikaite *et al.* | Invasive | 13 SNPs | UK | 2016 |
| ERS1594824 | Lynskey & Jauneikaite *et al.* | Invasive | 23 SNPs | UK | 2013 |
| ERS1594734 | Lynskey & Jauneikaite *et al.* | Invasive | 23 SNPs | UK | 2013 |
| ERS1594744 | Lynskey & Jauneikaite *et al.* | Invasive | 23 SNPs | UK | 2013 |
| ERS1594757 | Lynskey & Jauneikaite *et al.* | Invasive | 23 SNPs | UK | 2013 |
| ERS1594864 | Lynskey & Jauneikaite *et al.* | Invasive | 23 SNPs | UK | 2013 |
| ERS1594882 | Lynskey & Jauneikaite *et al.* | Invasive | 23 SNPs | UK | 2013 |
| ERS1594950 | Lynskey & Jauneikaite *et al.* | Invasive | 23 SNPs | UK | 2016 |

| | | | | | |
|---|---|---|---|---|---|
| ERS1595205 | Lynskey & Jauneikaite *et al.* | Invasive | 23 SNPs | UK | 2016 |
| ERS1594782 | Lynskey & Jauneikaite *et al.* | Invasive | M1global | UK | 2013 |
| ERS1594798 | Lynskey & Jauneikaite *et al.* | Invasive | M1global | UK | 2013 |
| ERS1594822 | Lynskey & Jauneikaite *et al.* | Invasive | M1global | UK | 2013 |
| ERS1594988 | Lynskey & Jauneikaite *et al.* | Invasive | M1global | UK | 2016 |
| ERS1595222 | Lynskey & Jauneikaite *et al.* | Invasive | M1global | UK | 2016 |
| ERS1595038 | Lynskey & Jauneikaite *et al.* | Invasive | M1global | UK | 2016 |
| ERS1594786 | Lynskey & Jauneikaite *et al.* | Invasive | M1uk | UK | 2013 |
| ERS1594722 | Lynskey & Jauneikaite *et al.* | Invasive | M1uk | UK | 2013 |
| ERS1594847 | Lynskey & Jauneikaite *et al.* | Invasive | M1uk | UK | 2013 |
| ERS1594947 | Lynskey & Jauneikaite *et al.* | Invasive | M1uk | UK | 2016 |
| ERS1595232 | Lynskey & Jauneikaite *et al.* | Invasive | M1uk | UK | 2016 |
| ERS1595131 | Lynskey & Jauneikaite *et al.* | Invasive | M1uk | UK | 2016 |
| ERS1020045 | Lynskey & Jauneikaite *et al.* | Non invasive | M1global | UK | 2009 |
| ERS1020472 | Lynskey & Jauneikaite *et al.* | Non invasive | M1global | UK | 2010 |
| ERS1020385 | Lynskey & Jauneikaite *et al.* | Non invasive | M1global | UK | 2009 |
| ERS1020523 | Lynskey & Jauneikaite *et al.* | Non invasive | M1global | UK | 2009 |
| ERS1020620 | Lynskey & Jauneikaite *et al.* | Non invasive | M1global | UK | 2015 |
| ERS1463088 | Lynskey & Jauneikaite *et al.* | Non invasive | M1uk | UK | 2016 |
| ERS1020174 | Lynskey & Jauneikaite *et al.* | Non invasive | M1uk | UK | 2012 |
| ERS1020714 | Lynskey & Jauneikaite *et al.* | Non invasive | M1uk | UK | 2015 |
| ERS1020508 | Lynskey & Jauneikaite *et al.* | Non invasive | M1uk | UK | 2015 |
| ERS1020090 | Lynskey & Jauneikaite *et al.* | Non invasive | M1uk | UK | 2011 |
| ERS1020076 | Lynskey & Jauneikaite *et al.* | Non invasive | M1global | UK | 2010 |
| ERS1020092 | Lynskey & Jauneikaite *et al.* | Non invasive | M1global | UK | 2010 |
| ERS1020107 | Lynskey & Jauneikaite *et al.* | Non invasive | M1global | UK | 2010 |
| ERS1020141 | Lynskey & Jauneikaite *et al.* | Non invasive | M1global | UK | 2010 |
| ERS1020171 | Lynskey & Jauneikaite *et al.* | Non invasive | 13 SNPs | UK | 2011 |
| ERS1020034 | Lynskey & Jauneikaite *et al.* | Non invasive | M1global | UK | 2011 |
| ERS1020036 | Lynskey & Jauneikaite *et al.* | Non invasive | M1global | UK | 2011 |
| ERS1020046 | Lynskey & Jauneikaite *et al.* | Non invasive | M1global | UK | 2011 |
| ERS1020050 | Lynskey & Jauneikaite *et al.* | Non invasive | 13 SNPs | UK | 2011 |
| ERS1020056 | Lynskey & Jauneikaite *et al.* | Non invasive | M1global | UK | 2011 |
| ERS1020079 | Lynskey & Jauneikaite *et al.* | Non invasive | M1global | UK | 2011 |
| ERS1020095 | Lynskey & Jauneikaite *et al.* | Non invasive | M1global | UK | 2011 |
| ERS1020102 | Lynskey & Jauneikaite *et al.* | Non invasive | M1global | UK | 2011 |
| ERS1020104 | Lynskey & Jauneikaite *et al.* | Non invasive | M1global | UK | 2011 |
| ERS1020128 | Lynskey & Jauneikaite *et al.* | Non invasive | M1global | UK | 2010 |
| ERS1020136 | Lynskey & Jauneikaite *et al.* | Non invasive | M1global | UK | 2012 |
| ERS1020152 | Lynskey & Jauneikaite *et al.* | Non invasive | M1global | UK | 2012 |
| ERS1020156 | Lynskey & Jauneikaite *et al.* | Non invasive | 23 SNPs | UK | 2012 |
| ERS1020158 | Lynskey & Jauneikaite *et al.* | Non invasive | M1global | UK | 2012 |
| ERS1020162 | Lynskey & Jauneikaite *et al.* | Non invasive | M1global | UK | 2010 |
| ERS1020199 | Lynskey & Jauneikaite *et al.* | Non invasive | M1global | UK | 2012 |
| ERS1020207 | Lynskey & Jauneikaite *et al.* | Non invasive | M1uk | UK | 2012 |
| ERS1020240 | Lynskey & Jauneikaite *et al.* | Non invasive | M1uk | UK | 2014 |
| ERS1020250 | Lynskey & Jauneikaite *et al.* | Non invasive | M1global | UK | 2014 |
| ERS1020275 | Lynskey & Jauneikaite *et al.* | Non invasive | M1uk | UK | 2014 |
| ERS1020277 | Lynskey & Jauneikaite *et al.* | Non invasive | M1global | UK | 2014 |
| ERS1020341 | Lynskey & Jauneikaite *et al.* | Non invasive | M1uk | UK | 2014 |
| ERS1020354 | Lynskey & Jauneikaite *et al.* | Non invasive | M1global | UK | 2014 |
| ERS1020358 | Lynskey & Jauneikaite *et al.* | Non invasive | M1uk | UK | 2014 |
| ERS1020374 | Lynskey & Jauneikaite *et al.* | Non invasive | M1global | UK | 2014 |
| ERS1020253 | Lynskey & Jauneikaite *et al.* | Non invasive | M1global | UK | 2014 |
| ERS1020361 | Lynskey & Jauneikaite *et al.* | Non invasive | M1global | UK | 2009 |
| ERS1020365 | Lynskey & Jauneikaite *et al.* | Non invasive | M1global | UK | 2009 |
| ERS1020393 | Lynskey & Jauneikaite *et al.* | Non invasive | M1global | UK | 2009 |
| ERS1020398 | Lynskey & Jauneikaite *et al.* | Non invasive | M1global | UK | 2009 |
| ERS1020429 | Lynskey & Jauneikaite *et al.* | Non invasive | M1global | UK | 2009 |
| ERS1020433 | Lynskey & Jauneikaite *et al.* | Non invasive | M1global | UK | 2009 |

| | | | | | |
|---|---|---|---|---|---|
| ERS1020441 | Lynskey & Jauneikaite *et al.* | Non invasive | M1global | UK | 2009 |
| ERS1020443 | Lynskey & Jauneikaite *et al.* | Non invasive | M1global | UK | 2009 |
| ERS1020465 | Lynskey & Jauneikaite *et al.* | Non invasive | 13 SNPs | UK | 2009 |
| ERS1020497 | Lynskey & Jauneikaite *et al.* | Non invasive | M1global | UK | 2009 |
| ERS1020539 | Lynskey & Jauneikaite *et al.* | Non invasive | M1global | UK | 2009 |
| ERS1020551 | Lynskey & Jauneikaite *et al.* | Non invasive | 13 SNPs | UK | 2009 |
| ERS1020555 | Lynskey & Jauneikaite *et al.* | Non invasive | M1global | UK | 2009 |
| ERS1020572 | Lynskey & Jauneikaite *et al.* | Non invasive | M1global | UK | 2009 |
| ERS1020576 | Lynskey & Jauneikaite *et al.* | Non invasive | M1global | UK | 2009 |
| ERS1020435 | Lynskey & Jauneikaite *et al.* | Non invasive | M1global | UK | 2010 |
| ERS1020460 | Lynskey & Jauneikaite *et al.* | Non invasive | M1global | UK | 2010 |
| ERS1020486 | Lynskey & Jauneikaite *et al.* | Non invasive | M1uk | UK | 2015 |
| ERS1020488 | Lynskey & Jauneikaite *et al.* | Non invasive | M1global | UK | 2015 |
| ERS1020502 | Lynskey & Jauneikaite *et al.* | Non invasive | M1uk | UK | 2015 |
| ERS1020514 | Lynskey & Jauneikaite *et al.* | Non invasive | M1global | UK | 2015 |
| ERS1020561 | Lynskey & Jauneikaite *et al.* | Non invasive | M1global | UK | 2015 |
| ERS1020602 | Lynskey & Jauneikaite *et al.* | Non invasive | M1global | UK | 2015 |
| ERS1020603 | Lynskey & Jauneikaite *et al.* | Non invasive | M1uk | UK | 2015 |
| ERS1020613 | Lynskey & Jauneikaite *et al.* | Non invasive | M1global | UK | 2015 |
| ERS1020634 | Lynskey & Jauneikaite *et al.* | Non invasive | M1uk | UK | 2015 |
| ERS1020638 | Lynskey & Jauneikaite *et al.* | Non invasive | M1global | UK | 2015 |
| ERS1020646 | Lynskey & Jauneikaite *et al.* | Non invasive | M1uk | UK | 2015 |
| ERS1020668 | Lynskey & Jauneikaite *et al.* | Non invasive | M1uk | UK | 2015 |
| ERS1020679 | Lynskey & Jauneikaite *et al.* | Non invasive | M1uk | UK | 2015 |
| ERS1020688 | Lynskey & Jauneikaite *et al.* | Non invasive | M1uk | UK | 2015 |
| ERS1020696 | Lynskey & Jauneikaite *et al.* | Non invasive | M1global | UK | 2015 |
| ERS1020702 | Lynskey & Jauneikaite *et al.* | Non invasive | M1uk | UK | 2015 |
| ERS1020703 | Lynskey & Jauneikaite *et al.* | Non invasive | M1uk | UK | 2015 |
| ERS1020705 | Lynskey & Jauneikaite *et al.* | Non invasive | M1uk | UK | 2015 |
| ERS1020715 | Lynskey & Jauneikaite *et al.* | Non invasive | M1global | UK | 2015 |
| ERS1020717 | Lynskey & Jauneikaite *et al.* | Non invasive | M1uk | UK | 2015 |
| ERS1020719 | Lynskey & Jauneikaite *et al.* | Non invasive | M1uk | UK | 2015 |
| ERS1020732 | Lynskey & Jauneikaite *et al.* | Non invasive | M1uk | UK | 2015 |
| ERS1020604 | Lynskey & Jauneikaite *et al.* | Non invasive | M1uk | UK | 2015 |
| ERS1020629 | Lynskey & Jauneikaite *et al.* | Non invasive | M1uk | UK | 2015 |
| ERS1020631 | Lynskey & Jauneikaite *et al.* | Non invasive | M1uk | UK | 2015 |
| ERS1020651 | Lynskey & Jauneikaite *et al.* | Non invasive | M1uk | UK | 2015 |
| ERS1020659 | Lynskey & Jauneikaite *et al.* | Non invasive | M1uk | UK | 2015 |
| ERS1020667 | Lynskey & Jauneikaite *et al.* | Non invasive | M1uk | UK | 2015 |
| ERS1020687 | Lynskey & Jauneikaite *et al.* | Non invasive | M1uk | UK | 2010 |
| ERS1020692 | Lynskey & Jauneikaite *et al.* | Non invasive | M1global | UK | 2012 |
| ERS1020694 | Lynskey & Jauneikaite *et al.* | Non invasive | M1uk | UK | 2012 |
| ERS1020701 | Lynskey & Jauneikaite *et al.* | Non invasive | M1global | UK | 2009 |
| ERS1463073 | Lynskey & Jauneikaite *et al.* | Non invasive | M1global | UK | 2016 |
| ERS1463078 | Lynskey & Jauneikaite *et al.* | Non invasive | M1uk | UK | 2016 |
| ERS1463079 | Lynskey & Jauneikaite *et al.* | Non invasive | M1global | UK | 2016 |
| ERS1463083 | Lynskey & Jauneikaite *et al.* | Non invasive | M1uk | UK | 2016 |
| ERS1463084 | Lynskey & Jauneikaite *et al.* | Non invasive | M1global | UK | 2016 |
| ERS1463085 | Lynskey & Jauneikaite *et al.* | Non invasive | M1uk | UK | 2016 |
| ERS1463086 | Lynskey & Jauneikaite *et al.* | Non invasive | M1uk | UK | 2016 |
| ERS1463087 | Lynskey & Jauneikaite *et al.* | Non invasive | M1uk | UK | 2016 |
| ERS1463089 | Lynskey & Jauneikaite *et al.* | Non invasive | M1uk | UK | 2016 |
| ERS1463090 | Lynskey & Jauneikaite *et al.* | Non invasive | M1uk | UK | 2016 |
| ERS1463091 | Lynskey & Jauneikaite *et al.* | Non invasive | M1uk | UK | 2016 |
| ERS1463092 | Lynskey & Jauneikaite *et al.* | Non invasive | M1uk | UK | 2016 |
| ERS1463093 | Lynskey & Jauneikaite *et al.* | Non invasive | M1uk | UK | 2016 |
| ERS1463095 | Lynskey & Jauneikaite *et al.* | Non invasive | M1global | UK | 2016 |
| ERS1463096 | Lynskey & Jauneikaite *et al.* | Non invasive | M1uk | UK | 2016 |
| ERS1463100 | Lynskey & Jauneikaite *et al.* | Non invasive | M1global | UK | 2016 |
| ERS1463101 | Lynskey & Jauneikaite *et al.* | Non invasive | M1uk | UK | 2016 |

| | | | | | |
|---|---|---|---|---|---|
| ERS1463102 | Lynskey & Jauneikaite *et al.* | Non invasive | M1global | UK | 2016 |
| ERS1463104 | Lynskey & Jauneikaite *et al.* | Non invasive | M1uk | UK | 2016 |
| ERS1463105 | Lynskey & Jauneikaite *et al.* | Non invasive | M1global | UK | 2016 |
| ERS1463106 | Lynskey & Jauneikaite *et al.* | Non invasive | M1global | UK | 2016 |
| ERS1463107 | Lynskey & Jauneikaite *et al.* | Non invasive | M1global | UK | 2016 |
| ERS1463108 | Lynskey & Jauneikaite *et al.* | Non invasive | M1global | UK | 2016 |
| ERS1463109 | Lynskey & Jauneikaite *et al.* | Non invasive | M1global | UK | 2016 |
| ERS1463110 | Lynskey & Jauneikaite *et al.* | Non invasive | M1global | UK | 2016 |
| ERS1463111 | Lynskey & Jauneikaite *et al.* | Non invasive | M1uk | UK | 2016 |
| ERS1463112 | Lynskey & Jauneikaite *et al.* | Non invasive | M1global | UK | 2016 |
| ERS1463113 | Lynskey & Jauneikaite *et al.* | Non invasive | M1uk | UK | 2016 |
| ERS1463114 | Lynskey & Jauneikaite *et al.* | Non invasive | M1global | UK | 2016 |
| ERS1463115 | Lynskey & Jauneikaite *et al.* | Non invasive | M1uk | UK | 2016 |
| ERS1463116 | Lynskey & Jauneikaite *et al.* | Non invasive | M1uk | UK | 2016 |
| ERS1463117 | Lynskey & Jauneikaite *et al.* | Non invasive | M1global | UK | 2016 |
| ERS1463118 | Lynskey & Jauneikaite *et al.* | Non invasive | M1uk | UK | 2016 |
| ERS1463119 | Lynskey & Jauneikaite *et al.* | Non invasive | M1uk | UK | 2016 |
| ERS1463120 | Lynskey & Jauneikaite *et al.* | Non invasive | M1global | UK | 2016 |
| ERS1463121 | Lynskey & Jauneikaite *et al.* | Non invasive | M1uk | UK | 2016 |
| ERS1463122 | Lynskey & Jauneikaite *et al.* | Non invasive | M1uk | UK | 2016 |
| ERS1463123 | Lynskey & Jauneikaite *et al.* | Non invasive | M1uk | UK | 2016 |
| ERS1463124 | Lynskey & Jauneikaite *et al.* | Non invasive | M1uk | UK | 2016 |
| ERS1463125 | Lynskey & Jauneikaite *et al.* | Non invasive | M1uk | UK | 2016 |
| ERS1463126 | Lynskey & Jauneikaite *et al.* | Non invasive | M1uk | UK | 2016 |
| ERS1463127 | Lynskey & Jauneikaite *et al.* | Non invasive | M1uk | UK | 2016 |
| ERS1463129 | Lynskey & Jauneikaite *et al.* | Non invasive | M1uk | UK | 2016 |
| ERS1463130 | Lynskey & Jauneikaite *et al.* | Non invasive | M1uk | UK | 2016 |
| ERS1463131 | Lynskey & Jauneikaite *et al.* | Non invasive | M1uk | UK | 2016 |
| ERS1463072 | Lynskey & Jauneikaite *et al.* | Non invasive | M1uk | UK | 2016 |
| ERS1463099 | Lynskey & Jauneikaite *et al.* | Non invasive | M1uk | UK | 2016 |
| ERS1463128 | Lynskey & Jauneikaite *et al.* | Non invasive | M1uk | UK | 2016 |
| ERS1463080 | Lynskey & Jauneikaite *et al.* | Non invasive | M1uk | UK | 2016 |
| ERS362081 | Turner *et al.* | Invasive | M1global | UK | 2006 |
| ERS362084 | Turner *et al.* | Invasive | M1global | UK | 2006 |
| ERS362085 | Turner *et al.* | Invasive | M1global | UK | 2006 |
| ERS362098 | Turner *et al.* | Invasive | M1global | UK | 2007 |
| ERS362108 | Turner *et al.* | Invasive | M1global | UK | 2011 |
| ERS362125 | Turner *et al.* | Invasive | M1global | UK | 2008 |
| ERS362128 | Turner *et al.* | Invasive | M1global | UK | 2009 |
| ERS362130 | Turner *et al.* | Invasive | M1global | UK | 2009 |
| ERS362131 | Turner *et al.* | Invasive | M1global | UK | 2009 |
| ERS362134 | Turner *et al.* | Invasive | M1global | UK | 2009 |
| ERS362138 | Turner *et al.* | Invasive | M1global | UK | 2010 |
| ERS361917 | Turner *et al.* | Invasive | M1global | UK | 2011 |
| ERS361974 | Turner *et al.* | Invasive | M1global | UK | 2007 |
| ERS361979 | Turner *et al.* | Invasive | M1global | UK | 2008 |
| ERS361984 | Turner *et al.* | Invasive | M1global | UK | 2008 |
| ERS361986 | Turner *et al.* | Invasive | M1global | UK | 2008 |
| ERS361987 | Turner *et al.* | Invasive | M1global | UK | 2008 |
| ERS361989 | Turner *et al.* | Invasive | M1global | UK | 2008 |
| ERS361995 | Turner *et al.* | Invasive | M1global | UK | 2009 |
| ERS362002 | Turner *et al.* | Invasive | M1global | UK | 2008 |
| ERS362004 | Turner *et al.* | Invasive | M1global | UK | 2009 |
| ERS362006 | Turner *et al.* | Invasive | M1global | UK | 2009 |
| ERS362102 | Turner *et al.* | Invasive | 19 SNPs | UK | 2007 |
| ERS362148 | Turner *et al.* | Invasive | M1global | UK | 2011 |
| ERS362149 | Turner *et al.* | Invasive | M1global | UK | 2011 |
| ERS362150 | Turner *et al.* | Invasive | M1global | UK | 2011 |
| ERS362141 | Turner *et al.* | Invasive | M1global | UK | 2010 |
| ERS362144 | Turner *et al.* | Invasive | M1global | UK | 2011 |

| | | | | | |
|---|---|---|---|---|---|
| ERS362145 | Turner *et al.* | Invasive | M1global | UK | 2011 |
| ERS362146 | Turner *et al.* | Invasive | M1uk | UK | 2011 |
| ERS362147 | Turner *et al.* | Invasive | M1uk | UK | 2011 |
| ERS362165 | Turner *et al.* | Invasive | M1global | UK | 2011 |
| ERS362031 | Turner *et al.* | Invasive | M1global | UK | 2005 |
| ERS362033 | Turner *et al.* | Invasive | M1global | UK | 2005 |
| ERS362040 | Turner *et al.* | Invasive | M1global | UK | 2005 |
| ERS362047 | Turner *et al.* | Invasive | M1global | UK | 2005 |
| ERS362051 | Turner *et al.* | Invasive | M1global | UK | 2006 |
| ERS362058 | Turner *et al.* | Invasive | M1global | UK | 2006 |
| ERS362063 | Turner *et al.* | Invasive | M1global | UK | 2006 |
| ERS362066 | Turner *et al.* | Invasive | M1global | UK | 2006 |
| ERS362067 | Turner *et al.* | Invasive | M1global | UK | 2006 |
| ERS362071 | Turner *et al.* | Invasive | 13 SNPs | UK | 2006 |
| ERS362110 | Turner *et al.* | Invasive | M1global | UK | 2003 |
| ERS362122 | Turner *et al.* | Invasive | M1global | UK | 2005 |
| ERS362029 | Turner *et al.* | Invasive | M1global | UK | 2005 |
| ERS361855 | Turner *et al.* | Invasive | M1global | UK | 2001 |
| ERS361867 | Turner *et al.* | Invasive | M1global | UK | 2002 |
| ERS361913 | Turner *et al.* | Invasive | M1global | UK | 2004 |
| ERS361937 | Turner *et al.* | Invasive | M1global | UK | 2004 |
| ERS361941 | Turner *et al.* | Invasive | M1global | UK | 2004 |
| ERS361950 | Turner *et al.* | Invasive | M1global | UK | 2004 |
| ERS361954 | Turner *et al.* | Invasive | M1global | UK | 2004 |
| ERS361960 | Turner *et al.* | Invasive | 13 SNPs | UK | 2005 |
| ERS361966 | Turner *et al.* | Invasive | M1global | UK | 2004 |
| ERS361929 | Turner *et al.* | Invasive | M1global | UK | 2004 |
| ERS361826 | Turner *et al.* | Invasive | M1global | UK | 2001 |
| ERS361827 | Turner *et al.* | Invasive | M1global | UK | 2001 |
| ERS361841 | Turner *et al.* | Invasive | M1global | UK | 2001 |
| ERS361865 | Turner *et al.* | Invasive | M1global | UK | 2002 |
| ERS362030 | Turner *et al.* | Invasive | M1global | UK | 2005 |
| ERS362032 | Turner *et al.* | Invasive | M1global | UK | 2005 |
| ERS362049 | Turner *et al.* | Invasive | M1global | UK | 2005 |
| ERS362078 | Turner *et al.* | Invasive | M1global | UK | 2006 |
| ERS379360 | Turner *et al.* | Invasive | M1global | UK | 2006 |

¶Non-invasive strains that were used in SpeA quantification (Supplementary figure 2) are highlighted by grey shading.

†Invasive strains representing four sublineages used for SpeA quantification are highlighted in blue shading

**Supplementary Table S3.** Components of chemically defined media (CDM) –

| COMPONENT¶ | CONSTITUENT | STOCK CONC | VOLUME IN 500mL CDM |
|---|---|---|---|
| Water | n/a | n/a | 350 |
| Iron | Fe(NO3)3.9H2O - 50mg | 1000X | 0.5 |
| | F2SO4.7H2O - 250mg | | |
| | made up to50mL ddH2O | | |
| Phosphate | K2HPO4 - 5g | 25X | 20 |
| | KH2PO4 - 25g | | |
| | NaH2PO4.H2O - 79.9g | | |
| | Na2HPO4.7H2O - 346.8g | | |
| | made up to 1L ddH2O | | |
| MgSO4.7H2O | MgSO4.7H2O - 87.5g | 500X | 1 |
| | made up to 250mL ddH2O | | |
| MnSO4.H2O | MnSO4.H2O - 1.25g | 1000X | 0.5 |
| | made up to 250mL ddH2O | | |
| Sodium acetate | NaC2H3O2.3H2O - 112.5g | 50X | 10 |
| | made up to 500mL ddH2O | | |
| Calcium chloride | CaCl2.2H2O - 338mg | 1000X | 0.5 |
| | made up to 50mL ddH2O | | |
| Sodium bicarbonate | NaHCO3 - 50g | 20X | 25 |
| | made up to 1L dd H2O | | |
| L-Cysteine HCl | Cysteine HCl - 16.25mg | 500X | 1 |
| | made up to 50mL ddH2O | | |
| Bases | 2N HCl - 50mL | 100X | 5 |
| | adenine - 0.5g | | |
| | guanine hydrochloride - 0.5g | | |
| | uracil - 0.5g | | |
| | ddH2O - 200ml | | |
| Vitamins | p-aminobenzoic acid - 10mg | 1000X | 0.5 |
| | biotin - 10mg | | |
| | folic acid - 40mg | | |
| | nicotinamide - 50mg | | |
| | b-nicotinamide adenine dinucleotide phosphate - 125mg | | |
| | pantothenate calcium salt - 100mg | | |
| | pyridoxal - 50mg | | |
| | pyridoxamine dihydrochloride - 50mg | | |
| | riboflavin - 100mg | | |
| | thiamine hydrochloride - 50mg | | |
| | vitamin B12 - 5mg | | |
| | dissolve in 50mL ddH2O | | |
| Amino acids | DL- alanine - 2.5g | 50X | 10 |

| | | | |
|---|---|---|---|
| | L-arginine - 2.5g | | |
| | L-aspartic acid - 2.5g | | |
| | L-asparagine - 2.5g | | |
| | L-cystine 1.25g | | |
| | L-glutamic acid - 2.5g | | |
| | L-glutamine - 5.0g | | |
| | Glycine - 2.5g | | |
| | L-histidine - 2.5g | | |
| | L-isoleucine - 2.5g | | |
| | L-leucine - 2.5g | | |
| | L-lysine - 2.5g | | |
| | L-methionine - 2.5g | | |
| | L-phenylalanine - 2.5g | | |
| | L-proline - 2.5g | | |
| | hydroxy-L-proline - 2.5g | | |
| | L-serine - 2.5g | | |
| | L-threonine - 5.0g | | |
| | L-tryptophan - 2.5g | | |
| | L-tyrosine - 2.5g | | |
| | L-valine - 2.5g | | |
| | made up to 500mL ddH2O | | |
| Carbon source | e.g., glucose | n/a | 5g |

¶adapted from van de Rijn and Kessler, *Infect Immun*, 27(2):444

**Supplementary Table S4**. Primers used for RT-qPCR

| GENE | PRIMER | SEQUENCE | PRODUCT (bp) |
|---|---|---|---|
| gldA | gldA_RTF | GCCTCAGATAATGAAATCAGCC | 121 |
| | gldA_RTR | CAAGTAAGTCAGCGATAGCC | |
| mipB | mipB_RTF | TAGGAGCACAAGCCATCAC | 100 |
| | mipB_RTR | TCCCAATCCTTGCCAAAATC | |
| pflD | pfld_RTF | CAAAAACGGCTAAACCAGAAC | 118 |
| | pfld_RTR | ATGAACCAGACAGATTGCAC | |
| ptsIIc | PTS_RTF | GCAAAACATCATCAAGCCAATC | 121 |
| | PTS_RTR | CCCAGCAATCAGGAAAAGAC | |
| glpF2 | glpF_RTF | GCTATGGCTTAGGAGTTATG | 120 |
| | glpF_RTR | AGACGTGAGCCCATGGGAAC | |
| speA | speA_Fwd | GAGGGGTAACAAATCATGAAGG | 94 |
| | speA_Rv | TCAAATGATAGGCTTTGGATACC | |
| proS | proS_Fwd | TGAGTTTATTATGAAAGACGGCTATAGTTTC | 93 |
| | proS_Rv | AATAGCTTCGTAAGCTTGACGATAATC | |
